# Local and biogeographical determinants of the diversity and structure of invertebrate communities in rot holes of ash, *Fraxinus excelsior*

**DOI:** 10.64898/2026.07.17.739164

**Authors:** Will Dawson, T. Hefin Jones, Jordan P. Cuff, Matthew Shepherd, Lynne Boddy

**Affiliations:** School of Biosciences, Cardiff University, Cardiff, United Kingdom, CF10 3AX; School of Natural and Environmental Sciences, Newcastle University, Newcastle-upon-Tyne, United Kingdom; Fera Science Ltd., York Biotech Campus, York, United Kingdom; Natural England, Horizon House, Deanery Road, Bristol, United Kingdom

**Keywords:** saproxylic, ancient woodland, veteran trees, tree hollows, deadwood, dieback

## Abstract

1. Veteran trees are biodiversity hotspots, providing a range of microhabitats, including tree hollows or rot holes in trunks, which support a large diversity of invertebrates including threatened species. Each rot hole is a unique microcosm, with variable biotic and abiotic characteristics enabling their use by many invertebrate taxa.
2. Most previous studies describing the invertebrate communities of rot holes and the factors determining their structure focus on a narrow range of invertebrate taxa, and many host tree species remain unexplored. European ash (*Fraxinus excelsior* L.) is one of Europe’s most threatened deciduous tree species due to the spread of ash dieback disease, but the potential impact of these losses on the invertebrate communities that inhabit ash is poorly understood.
3. Invertebrates were sampled and identified to family level from 14 sites across Wales and western England, and major physical characteristics of each rot hole and sample site were assessed (e.g., habitat type, altitude, maximum site temperature).
4. Ash rot hole invertebrate communities were highly diverse with a range of threatened taxa, including some undescribed or new to the region. There was large variation in both measured rot hole characteristics and the invertebrate communities between individual rot holes, sites and habitats. Both tree-specific and biogeographical factors significantly influenced invertebrate diversity and community structure, suggesting that sustaining the heterogeneity of ash rot holes across a range of locations is required to support saproxylic invertebrate diversity.
5. With increasing ash veteran tree vulnerability, sustaining these vital habitats and their associated invertebrate communities is a conservation imperative. This study has demonstrated that considering heterogeneity in both habitat characteristics and landscape context will be crucial when conserving these communities.

## Introduction

As they age, trees increasingly provide unique tree-related microhabitats (TreMs; Larrieu *et al*., 2018) which are utilised by many organisms. Veteran trees offer a high diversity of TreMs (Kozák *et al*., 2023), supporting different vertebrate and invertebrate communities (Spina *et al*., 2024; Majdi *et al*., 2025) vital for ecosystem functioning and biodiversity (Baeten *et al*., 2013; Lonsdale *et al*., 2013; Nakládal *et al*., 2022).

Among the most ecologically important and studied TreMs are tree hollows or rot holes. These are cavities within the trunk or larger limbs of a tree, usually formed by fungal decomposition of heartwood (Ranius *et al*., 2024). Rot holes are more abundant and structurally diverse in veteran trees than in younger trees (Wesołowski & Martin, 2018; Pilskog *et al*., 2020). Each rot hole is a microcosm with unique conditions, structure and substrate for inhabiting organisms (Quinto *et al*., 2014; Micó *et al*., 2015; Micó, 2018). The sheltered nature of rot holes enables the communities they contain to persist for some time following the death of the host tree (Stahlheber *et al*., 2015; Lindman *et al*., 2022; 2023).

While rot holes are utilised by over 2000 vertebrate species worldwide (e.g., birds nesting, bats roosting; Bryant *et al*., 2012; Stokland *et al*., 2012; Manning *et al*., 2013; Yatsiuk & Wesołowski, 2020), the animal communities are numerically dominated by invertebrates, some of which spend their entire life within a single rot hole (Hedin *et al*., 2008). In the United Kingdom alone, over 1800 invertebrate species depend on this type of rotting wood habitat, including numerous rare species (Alexander, 2002). Rot holes are clearly keystone habitats in forests (Müller *et al*., 2014), with approximately a third of all temperate forest invertebrates potentially saproxylic (Müller *et al*., 2008). Nevertheless, rot hole invertebrate species richness and abundances, as with insects in general (Seibold *et al*., 2019; Wagner, 2020), are declining globally (Manning *et al*., 2013; Seibold *et al*., 2015; Cálix *et al*., 2018; Schmidl, 2021; Quinto *et al*., 2023).

The long-term survival of rot hole invertebrate species and communities is ultimately dependent on host tree survival. Nevertheless, the number of veteran trees and their associated heart-rot habitat is severely declining globally (Dubois *et al*., 2009; Lindenmayer *et al*., 2014; Wesołowski *et al*., 2018), due largely to factors including the timber industry, forest management policies, urbanisation and disease (Boddy *et al*., 2017; Yatsiuk & Wesołowski, 2020). Though rate of decline is hard to quantify, in Norway net losses have been estimated as 1.2 % annually since 2011 (Jacobsen *et al*., 2023). There are more veteran trees in the UK than most of Europe combined, but losses have left a large age gap between the cohort with hollows and younger trees without, and veteran tree populations have become fragmented (Bengtsson *et al*., 2015; Nolan *et al*., 2020, 2022). Whilst some rot hole invertebrates can initially survive in fragmented veteran woodland populations (Davies *et al*., 2008), many have poor dispersal abilities, rendering them vulnerable (Ranius, 2002; 2006).

European ash (*Fraxinus excelsior* L.) is one of the most common tree species in Europe (Hladká, 2006; Maskell *et al*., 2013) and a vital keystone species in temperate deciduous woodlands (Littlewood et al., 2015; Hultberg *et al*., 2020). Ash dieback disease has decimated populations and threatens the future of ash TreMs and the organisms that depend on them (Coker *et al*., 2019; Combes et al., 2024). Previous studies on rot hole invertebrate communities have focused on either saproxylic beetles (e.g., Micó *et al*., 2013; Schauer *et al*., 2018; Henneberg *et al*., 2021; Quinto *et al*., 2023) or on non-ash tree species (e.g., Taylor & Ranius, 2014; Cuff *et al*., 2021b; Schauer *et al*., 2025), with only one such study including saproxylic arthropods in ash. Even then the latter study was carried out only in a single national park in Spain and did not include the entire mesofauna community (Quinto *et al*., 2014). There is a dearth of information on factors that determine community structure and diversity in ash rot holes, despite such knowledge likely being vital for the prioritisation of management and conservation.

Here the community structure of meso- and macro-invertebrates in ash rot holes, the effects of rot hole characteristics, and the climatic and geographical drivers at tree-, forest-, and biogeographic-scales were investigated. It was hypothesised that ash rot holes contain highly diverse invertebrate communities that (i) are structurally complex, with many under-recorded taxa; (ii) relate to tree-specific characteristics (e.g., characteristics of the host tree or wood rot within the rot hole); (iii) differ both locally and regionally, and (iv) are influenced by wider biogeographical factors due to different taxa being adapted for different habitats and climatic regimes (e.g., taxa more vulnerable to desiccation being influenced by rainfall and temperature).

## Methods

### Study Sites and Sample Collection

Individual rot holes (*n* = 110) were sampled between late September and November 2023, before the first winter frost. Fourteen sites were selected, using personal knowledge and the Ancient Tree Inventory (2023), across Wales and western England in four broad habitat types (low intensity local pasture farmland, parkland, urban woodland, non-urban woodland; Figure 1; Table 1). Samples were taken across six nationally spatial regions, with region referring to the spatial descriptors rather than distinct biogeographical regions. Habitats were categorised based on the dominant habitat type, with urban woodland distinguished from non-urban woodland by the presence of urbanised areas surrounding most of the woodland. Sample size within sites was determined by the availability, accessibility and suitability of rot holes.

**Figure 1.**
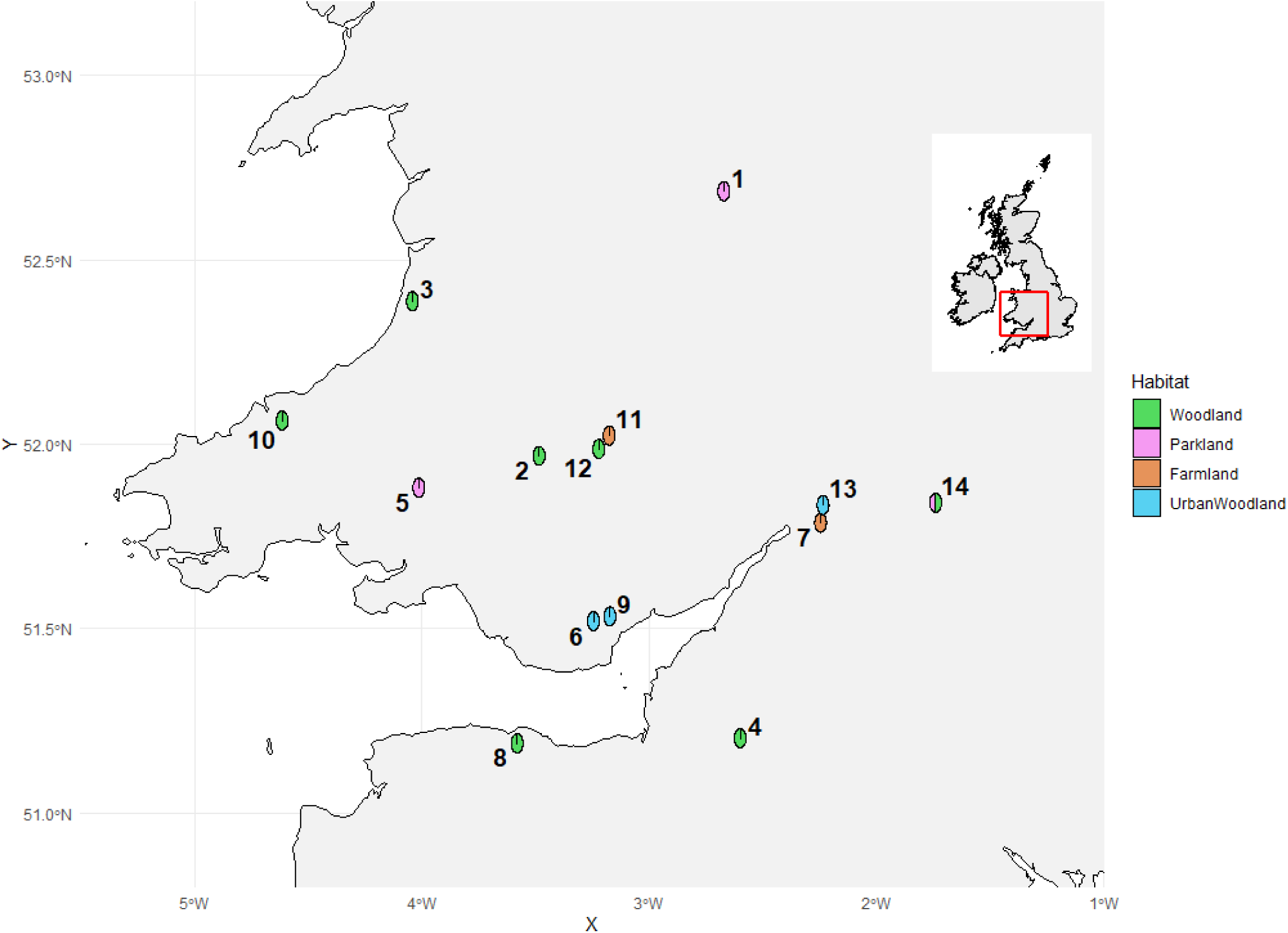
Sample sites in Wales and western England. The number adjacent to each point corresponds to each sample site. Site numbers and further information are given in Table 1.

**Table 1.**
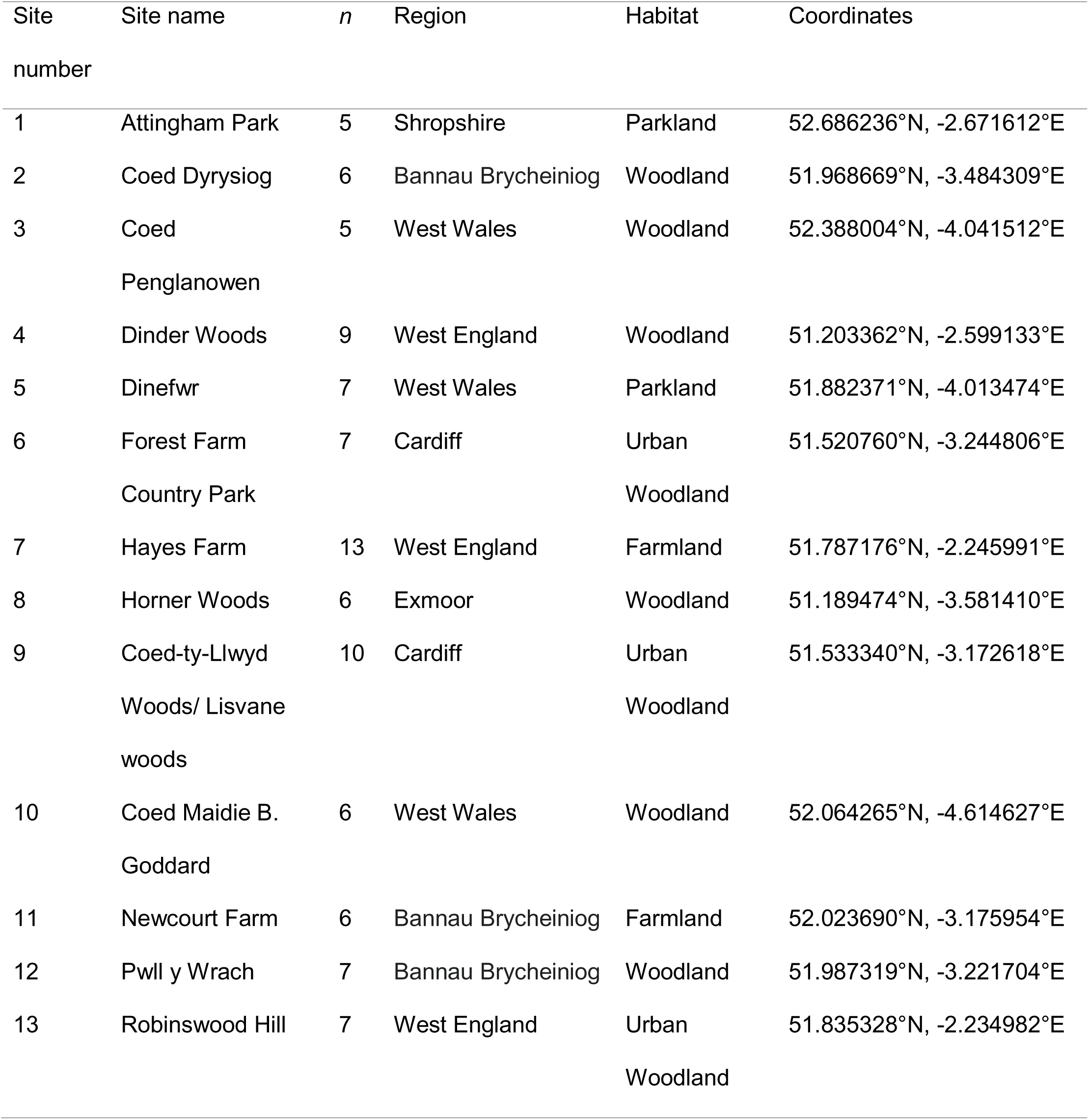
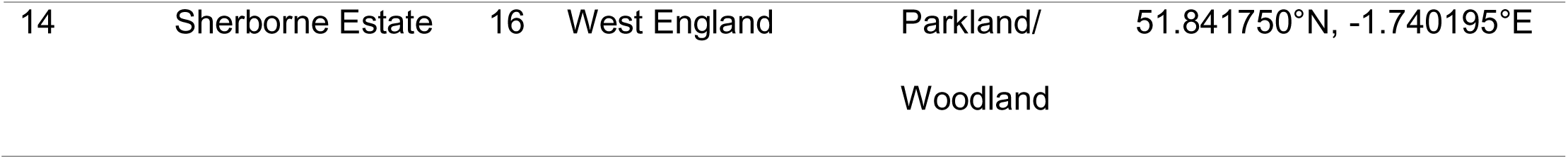
Site details for each of the locations from which rot holes were sampled. The number of samples for each site (*n*) is given alongside the region and broad habitat type.

| Site number | Site name | *n* | Region | Habitat | Coordinates |
| --- | --- | --- | --- | --- | --- |
| 1 | Attingham Park | 5 | Shropshire | Parkland | 52.686236°N, -2.671612°E |
| 2 | Coed Dyrysiog | 6 | Bannau Brycheiniog | Woodland | 51.968669°N, -3.484309°E |
| 3 | Coed Penglanowen | 5 | West Wales | Woodland | 52.388004°N, -4.041512°E |
| 4 | Dinder Woods | 9 | West England | Woodland | 51.203362°N, -2.599133°E |
| 5 | Dinefwr | 7 | West Wales | Parkland | 51.882371°N, -4.013474°E |
| 6 | Forest Farm Country Park | 7 | Cardiff | Urban Woodland | 51.520760°N, -3.244806°E |
| 7 | Hayes Farm | 13 | West England | Farmland | 51.787176°N, -2.245991°E |
| 8 | Horner Woods | 6 | Exmoor | Woodland | 51.189474°N, -3.581410°E |
| 9 | Coed-ty-Llwyd Woods/ Lisvane woods | 10 | Cardiff | Urban Woodland | 51.533340°N, -3.172618°E |
| 10 | Coed Maidie B. Goddard | 6 | West Wales | Woodland | 52.064265°N, -4.614627°E |
| 11 | Newcourt Farm | 6 | Bannau Brycheiniog | Farmland | 52.023690°N, -3.175954°E |
| 12 | Pwll y Wrach | 7 | Bannau Brycheiniog | Woodland | 51.987319°N, -3.221704°E |
| 13 | Robinswood Hill | 7 | West England | Urban Woodland | 51.835328°N, -2.234982°E |
| 14 | Sherborne Estate | 16 | West England | Parkland/<br>Woodland | 51.841750°N, -1.740195°E |

Wearing gloves, a grab sample (∼ 1 L; never more than 50 % of available rot) was taken from rot holes at the base of standing trees and transferred to a zip-lock bag with large air head space for transit. Prior to analysis, samples were kept at 4 °C, for no longer than 48 hours. The following tree characteristics were measured: the height of the rot hole above ground level, the area of opening (width * height), and the circumference of the trunk at breast height (1.3 m). Opening aspect was determined with a compass and recorded in degrees. Climatic data (maximum and minimum temperatures, and rainfall) were obtained from the nearest weather station to each site.

### Sorting and Rot Characteristics

In the laboratory, rot samples were emptied into foil trays. A ∼50 g (wet weight) subsample of rot was removed for rot characteristic analysis: fresh volume (*V*; cm^3^ determined in a measuring cylinder); wet mass (*m*_wet_; g); oven-dried weight after 72 h at ∼70 °C (*m*_dry_; g). These measurements enabled calculation of density (*d*) and gravimetric water content (*u*):

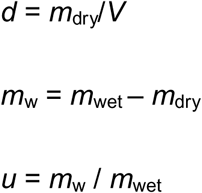

Note that care must be taken when interpreting percentage gravimetric water content as this value will differ for the same unit volume of water in rot of different density (Boddy, 1986) but is commonly used in studies of this type (e.g., Cuff *et al*., 2021b).

Clumped rot was broken apart by hand and any macroinvertebrates found were placed in 70 % ethanol. Invertebrates were extracted by placing the remaining rot material in Tullgren funnels (Burkard Scientific, Uxbridge, UK) for 72 h. All invertebrates extracted were collected into and stored in 70 % ethanol.

Invertebrate identification was undertaken under an Olympus SZX7 stereomicroscope with dichotomous taxonomic keys (Table S1). There are limited identification resources for many saproxylic taxa, and identification of larval stages, frequently prevalent in rot holes, is difficult (Gossner & Petermann, 2022; Köhler *et al*., 2022; Sire *et al*., 2025); hence, identification was made to family level to reduce bias when quantifying the diversity of different taxonomic groups. Nematodes and rotifers were not included in the study as Tullgren funnel dry extractions is biased against them (Schmelz & Collado, 2010). The phoretic deutonymph stage of Astigmatina (Sarcoptiformes: Oribatida) mites (hypopi) were recorded as a single group due to the difficulty involved in achieving finer taxonomic resolution.

### Statistical Analysis

Statistical analyses were performed in R version 4.4.0 (R Core Team, 2024). Independent variables were divided into tree-scale and biogeographical variables to improve model performance. Tree-scale factors included tree circumference, height of the rot hole on the tree, rot hole opening area, opening direction, and rot properties, whereas biogeographical factors were the sample site, habitat, region, maximum temperature, minimum temperature, rainfall and altitude. Direction was split into northness (cosine of direction angle) and eastness (sine of direction) to calculate comparable scores for analysis. Abundances were converted into relative frequencies (i.e., individual species abundances divided by total sample abundance) to standardise across communities. Rot hole communities were described using abundance, richness, and diversity. Diversity was investigated at three scales (Hulbert, 1971): each individual rot hole (α), within each woodland site (β) and across all woodland sites (γ). The α-diversity was calculated as Simpson’s diversity index (Hulbert, 1971). Kruskal-Wallis rank sum tests were used to investigate the variation in α-diversity across tree-scale and biogeographical factors (Ruxton & Beauchomp, 2008). For β-diversity, Bray-Curtis indices in the ‘betapart’ package (Baselga *et al*., 2018) were used to identify community nestedness and turnover (Baselga, 2017). Sampling completeness for the overall dataset, each habitat type, regions and sites sampled, and for individual samples were calculated using extrapolation and rarefaction of diversity with ‘iNEXT’ (Chao *et al*., 2014; Hsieh et al., 2016). Visualisation of the invertebrate community taxonomy was achieved using the ‘Metacoder’ package (Foster *et al*., 2017)

The interaction between different explanatory variables was assessed via correlation analysis using ‘corrplot’ (Wei *et al*., 2017). Principal component analysis (PCA; Abdi & Williams, 2010) was used to determine the tree-specific and biogeographical variables that relate to invertebrate community structure. The two most significant components calculated from PCA were then compared to community structure and nestedness using multivariate generalised linear models in ‘mvabund’ (M-GLMs; Warton *et al*., 2012). The influence of tree-specific and biogeographical factors, and their interactions upon both communities and individual taxonomic groups, were also determined using M-GLMs with negative binomial error family and Monte Carlo resampling.

Finally, co-occurrence of invertebrate taxa was assessed by comparing presence-absence matrices to null models with the ‘co-occur’ package (Griffith et al., 2016). This calculates the probability that two species would co-occur more or less frequently compared to an independent distribution across sites (Veech, 2013). One thousand random matrices were compared to the presence-absence matrices to identify non-random co-occurrences.

## Results

### Invertebrate community structure

From the sampled 110 rot holes, the invertebrates collected (45,698 individuals) were associated with 259 families across 41 orders and 11 classes (Figure 2, Table S2). The most abundant family was Isotomidae (Entomobryomorpha; *n* = 7,116), found in 92 % of samples (*n* = 101). Thirty families were represented by a single individual, and 37 families were found in one single rot hole (Table S2). While taxa were identified to family level, several notable specimens were identified further, including many individuals in the family Ichthystomatogasteridae (Mesostigmata), the first UK record of *Sejus sejiformis* (Balogh 1938; Mesostigmata: Sejidae), and a potential new species of *Puliciphora* (Diptera: Phoridae; unpublished data). The mean diversity within individual rot holes (alpha diversity) was α = 0.8692 (± 0.0942), and community composition varied between sites, regions and habitats (Figure 3; Figure S1). Similarly, alpha diversity significantly varied across sites (Kruskal-Wallis *X*^2^_13_ = 28.137, *p* = 0.009), region (Kruskal-Wallis *X*^2^_5_ = 13.866, *p* = 0.016), and habitat (Kruskal-Wallis *X*^2^_3_ = 13.038, *p* = 0.0046). This difference in diversity across sites (β = 0.981) was almost entirely driven by community turnover (β = 0.981) rather than nestedness (β = 9.955e^-13^). Sampling completeness across the study was estimated to be 99.93 %, with a high sample coverage across the four habitat types (> 99.5 % for each habitat).

**Figure 2.**
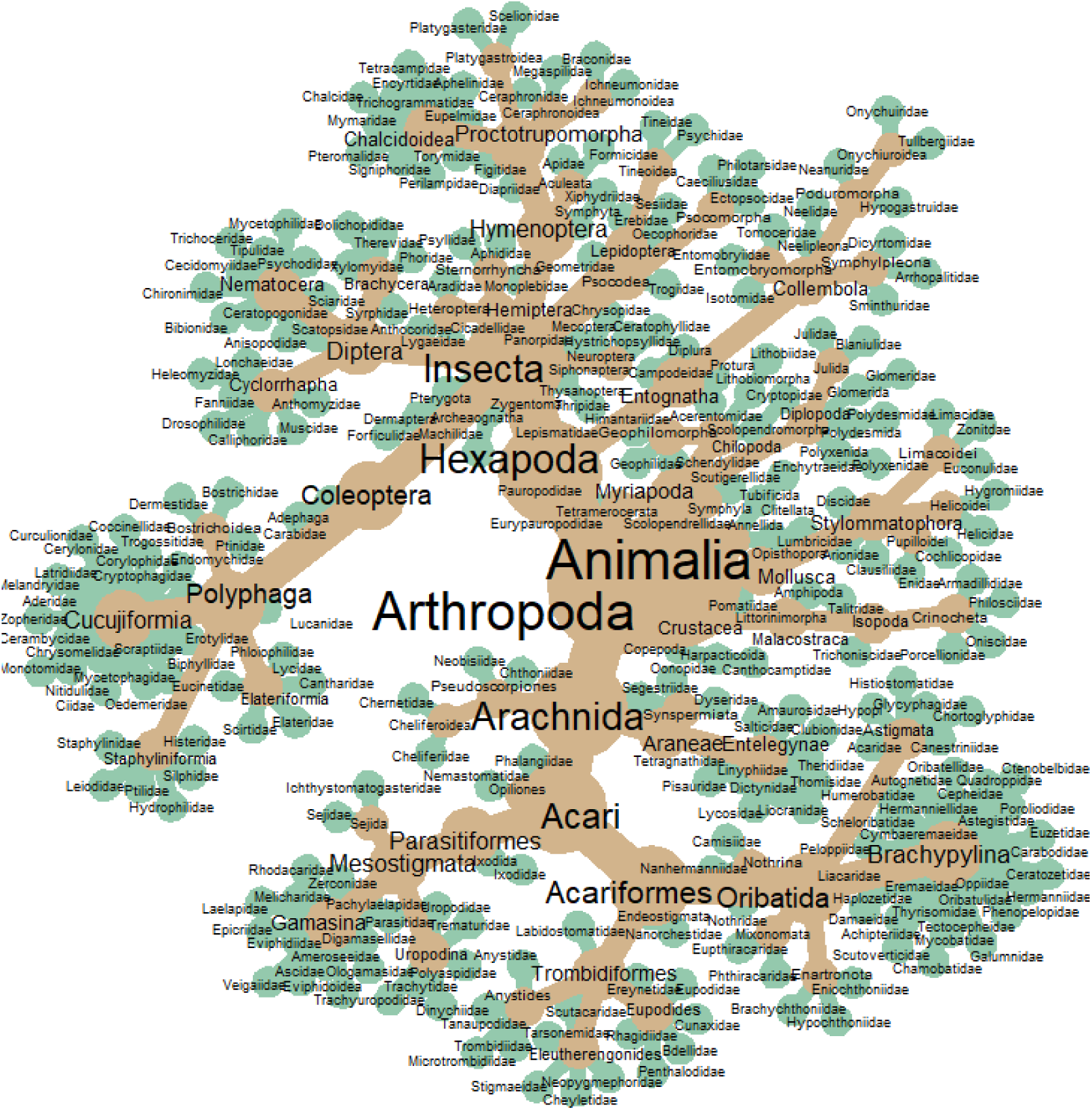
Metacoder plot of taxonomic diversity of invertebrates from ash rot holes. Families are at end nodes, and higher ranks are provided where possible to distinguish between coarser taxonomic groups.

**Figure 3.**
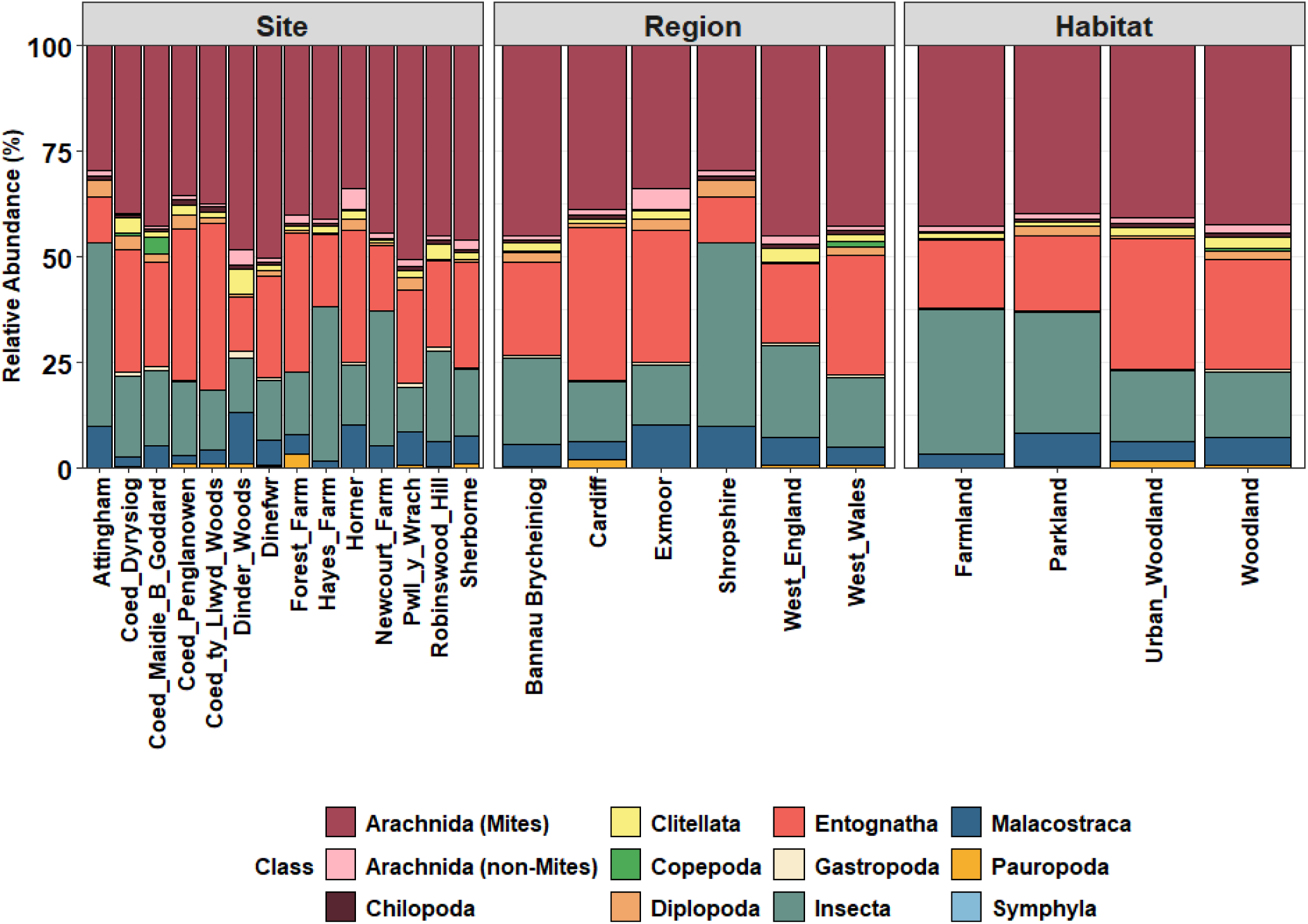
Relative abundance of invertebrate classes in ash rot holes for each sample site, region and habitat. Arachnida was split into subclass Acari and other Arachnida due to the high abundance of mites masking contributions by non-mite arachnids within the same samples.

### Influence of tree-scale variation

Tree-scale variables varied considerably between rot holes (Figure 4). Whilst the direction (both northness and eastness) and height of the rot holes did not differ significantly across sample sites, all other rot hole and tree-scale properties did (Table S3). Tree circumference (Kruskal-Wallis *X*^2^_3_ = 35.075, *p* < 0.001), rot water content (Kruskal-Wallis *X*^2^_3_ = 14.065, *p* = 0.003) and rot density (Kruskal-Wallis *X*^2^_3_ = 13.147, *p* = 0.004) varied significantly between habitat types but only rot density (Kruskal-Wallis *X*^2^_5_ = 18.422, *p* = 0.002) significantly differed between regions. Larger trees had significantly larger rot hole openings (*R*^2^ = 0.43, *p* < 0.001) and rot was drier than in smaller trees (*R*^2^ = -0.36, *p* < 0.001). While rot water content was positively correlated with the height of the rot hole (*R*^2^ = 0.25, *p* = 0.009), it was negatively related to the density of the wood rot inside (*R*^2^ = -0.75, *p* < 0.001). The northness of the opening direction was positively correlated with both the height of the rot hole on the tree (*R*^2^ = 0.20, *p* = 0.032) and rot water content (*R*^2^ = 0.20, p = 0.038) but negatively with the opening area (*R*^2^ = -0.26, p = 0.007; Figure 5).

**Figure 4.**
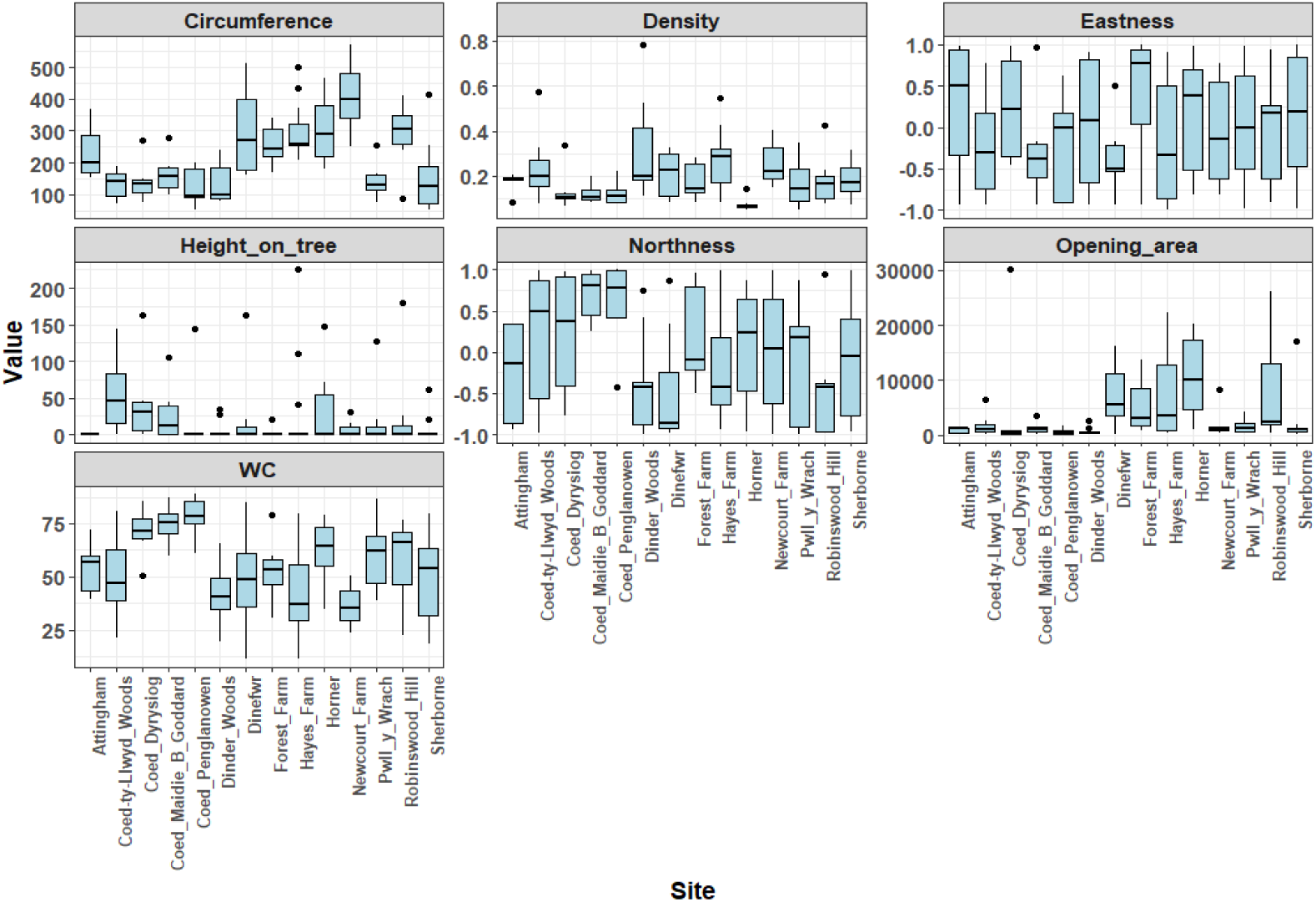
Variation in tree-scale variables across sample sites. Variables were altitude, host tree circumference, height of the rot hole on the tree, the rot hole opening area, as well as the wood rot density and gravimetric water content (WC).

**Figure 5.**
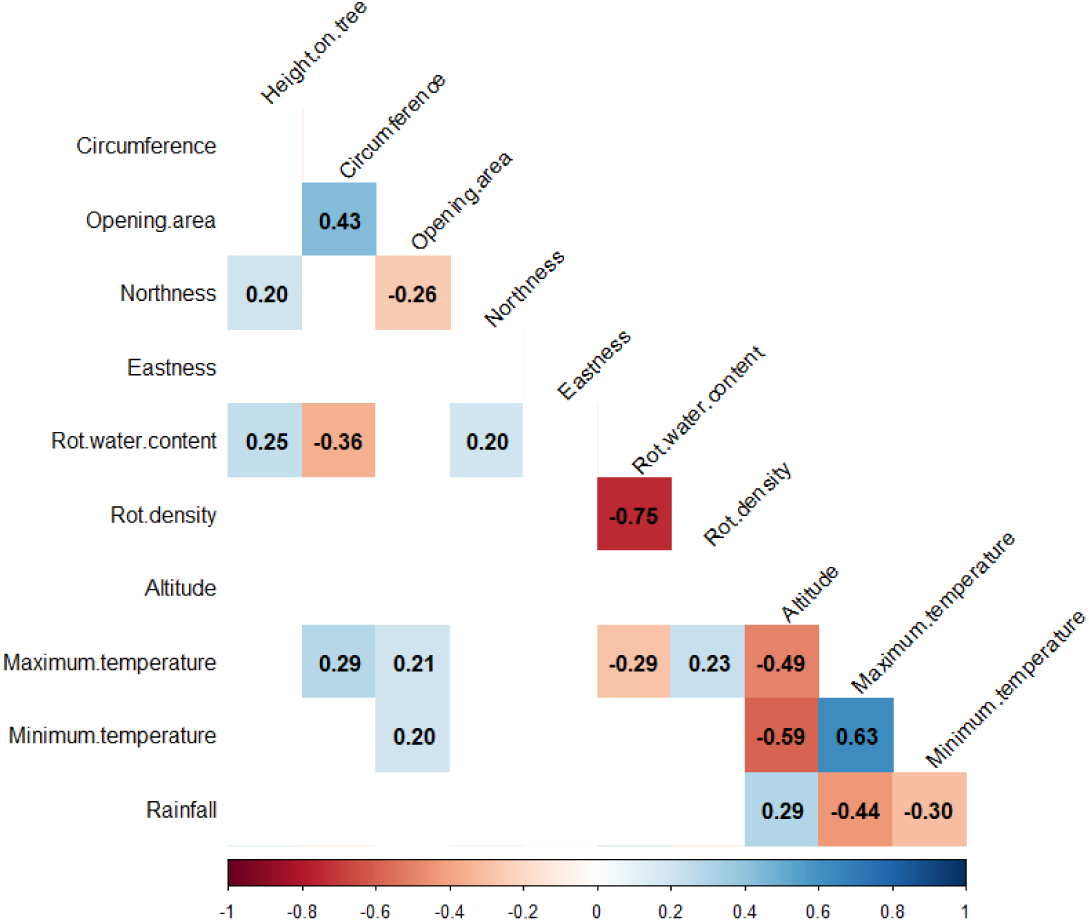
Correlogram of the relationships between different rot hole variables. Only significant correlations are displayed, with red denoting a negative correlation and blue a positive correlation.

The most significant two PCs relating to tree-scale factors explained 50.60 % of the invertebrate community variation (Figure 6). PC1 explained 30.81 % of the variation and correlated positively with wood rot water content (0.584) but negatively with rot density (-0.515), suggesting that the main drivers of invertebrate community structure were related to wood rot characteristics. PC2 explained 19.79 % of variation and positively correlated with rot hole opening size (0.696). The nestedness of the rot hole communities was not related to either of the tree-scale PC’s (*R*^2^ = 0.013, *F*_2,106_ = 0.71, *p* = 0.494).

**Figure 6.**
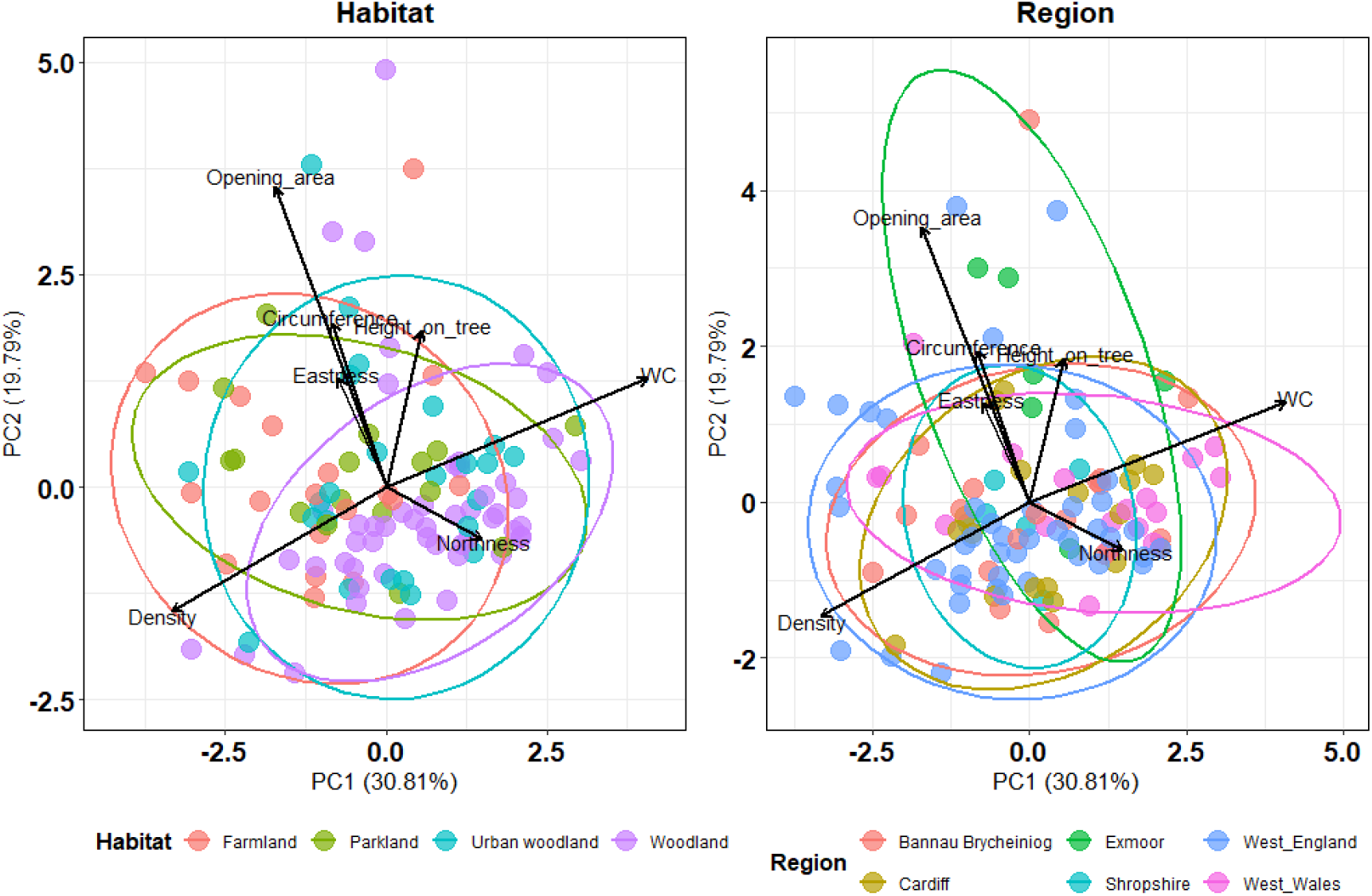
PCA results for tree-scale variables measured across ash rot holes. Arrow length shows the relative strength of correlation between the variables and PCs.

Invertebrate community composition was significantly related to all tree-scale variables (p < 0.05), including interactions between variables, except for the interactions of rot density with tree circumference, northness of the opening, and water content (Table 2). Fifteen taxa were significantly related to specific tree-scale variables or their interactions (Table S4, univariate GLMs): six Diptera (Syrphidae, Bibionidae, Anisopodidae, Ceratopogonidae, Muscidae and Scatopsidae), two Acari (Uropodidae and Ceratozetidae), and one each of Araneae (Lycosidae), Coleoptera (Lucanidae), Hymenoptera (Perilampidae), Isopoda (Armadillididae), Lepidoptera (Oecophoridae), Collembola (Hypogastruidae), and Diplopoda (Polyxenidae) families.

**Table 2.** Influence of tree-scale variables and their interactions on invertebrate community composition. The summary of MGLM outputs for the investigation of the influence of tree-scale variables (both individually and their interactions with each other) upon invertebrate community composition is provided with the deviation (*Dev*), degrees of freedom (*d.f*.) and significance (*p*).

| Variable/Interaction | Dev | d.f | p |
| --- | --- | --- | --- |
| Height on tree | 364.9247 | 108 | 0.001 |
| Circumference | 412.3468 | 107 | 0.001 |
| Opening area | 379.8795 | 106 | 0.001 |
| Eastness | 367.2455 | 105 | 0.001 |
| Northness | 469.0033 | 104 | 0.005 |
| Water content | 310.5037 | 103 | 0.002 |
| Density | 339.9526 | 102 | 0.001 |
| Height on tree:Circumference | 236.7221 | 101 | 0.001 |
| Height on tree:Opening area | 257.7993 | 100 | 0.004 |
| Height on tree:Eastness | 380.0893 | 99 | 0.001 |
| Height on tree:Northness | 298.1863 | 98 | 0.003 |
| Height on tree:Water content | 297.9826 | 97 | 0.002 |
| Height on tree:Density | 275.6948 | 96 | 0.005 |
| Circumference:Opening area | 349.7316 | 95 | 0.002 |
| Circumference:Angle | 343.0437 | 94 | 0.002 |
| Circumference:Water content | 281.5598 | 93 | 0.004 |
| Circumference:Density | 501.1823 | 92 | 0.360 |
| Opening area:Eastness | 389.9477 | 91 | 0.003 |
| Opening area:Northness | 4196.044 | 90 | 0.002 |
| Opening area:Water content | 298.0475 | 89 | 0.014 |
| Opening area:Density | 3873.393 | 88 | 0.002 |
| Eastness:Water content | 364.4701 | 87 | 0.001 |
| Northness:Water content | 5109.096 | 86 | 0.005 |
| Eastness:Density | 7351.435 | 85 | 0.022 |
| Northness:Density | 19798.32 | 84 | 0.262 |
| Water content:Density | 924.4749 | 83 | 0.652 |

### Influence of biogeographical variation

Invertebrate community composition from individual rot holes did not show clustering at sample site, region or habitat level. Alpha diversity significantly differed between habitats (Kruskal-Wallis *X*^2^_3_ = 13.038; *p* = 0.0046), but not between sites (Kruskal-Wallis *X*^2^_13_ = 28.137; *p* = 0.009) nor regions (Kruskal-Wallis *X*^2^_5_ = 13.866; *p* = 0.016). Parkland samples were the most biodiverse (α = 0.895 ± 0.048), followed by woodland (α = 0.893 ± 0.077) and urban woodland (0.835 ± 0.106), with farmland samples being the least diverse (α = 0.829 ± 0.124).

Rot hole elevation varied from 34 m to 239 m above sea level and varied significantly across sample sites (Kruskal-Wallis *X*^2^_13_ = 97.044, *p* < 0.001), regions (Kruskal-Wallis *X*^2^_5_ = 91,083, *p* < 0.001) and habitats (Kruskal-Wallis *X*^2^_13_ = 97.044, *p* < 0.001; Table S3). Altitude was negatively correlated with the maximum (*R*^2^ = - 0.49, p <0.001) and minimum (*R*^2^ = -0.59, p < 0.001) temperatures, but positively with rainfall (*R*^2^ = 0.29, p < 0.001; Figure 4). Warmer sites had less rainfall and larger trees with larger openings to their rot holes than cooler sites.

The first two PCs explained 69.85 % of variation in biogeographical factors, with PC1 explaining 43.47 % of variation and being negatively correlated with maximum temperature (-0.539), minimum temperature (-0.521) and positively correlated with altitude (0.427). PC2 explained 26.37 % of variation and was positively correlated to the latitude of the rot hole (0.761) and negatively to rainfall (- 0.448). As with tree-scale variables, the nestedness of the rot hole communities was not related to either biogeographical PC (*R*^2^ = 0.041, *F*_2,106_ = 2.262, *p* = 0.109).

Invertebrate community composition was significantly related to biogeographical variables (Table 3). Community composition differed significantly between sample sites (MGLM: *Dev*_96_ = 4565.309, *p* = 0.001), habitats (*Dev*_93_ = 267.243, *p* = 0.001) but not regions (*Dev*_89_ = 0.010, *p* = 0.819). Of the other biogeographic variables, only altitude (*Dev*_88_ = 414.235, *p* = 0.001) and its interactions with sample site (*Dev*_44_ = 1039.524, *p* = 0.001), habitat (*Dev*_77_ = 955.212, *p* = 0.001) and region (*Dev*_66_ = 975.378, *p* = 0.001) were significantly related to community composition. Seventeen individual families were significantly influenced by specific variables and their interactions (Table S5): six Acari (families Glycyphagidae, Ichthystomatogasteridae, Polyaspidiidae, Euzetidae, Scutacaridae and grouped hypopi), four Diptera (Tipulidae, Muscidae, Psychodidae and Scatopsidae), one Coleoptera (Ptiliidae), one Lepidoptera (Tineidae), one Diplopoda (Julidae), one Collembola (Onychuiridae), one Protura (Acerentomidae), one Thysanoptera (Thripidae), and one slug (Arionidae) family.

**Table 3.** Influence of biogeographical variables and their interactions on community structure. The summary of MGLMs investigating the influence of biogeographical variables (both individually and their interactions with each other) upon invertebrate community composition is provided with the deviation (*Dev*), degrees of freedom (*d.f*.) and significance (*p*) provided.

| Variable | Dev | d.f. | p |
| --- | --- | --- | --- |
| Sample site | 4565.309 | 96 | 0.001 |
| Habitat | 267.2433 | 93 | 0.001 |
| Region | 0.009962 | 89 | 0.819 |
| Altitude | 414.2353 | 88 | 0.001 |
| Maximum temperature | 0.095504 | 87 | 0.866 |
| Minimum temperature | 0.023125 | 86 | 0.711 |
| Rainfall | 0.234516 | 85 | 0.783 |
| Sample site:Habitat | 0.028816 | 82 | 0.989 |
| Habitat:Region | 0.00394 | 80 | 0.989 |
| Habitat:Altitude | 955.212 | 77 | 0.001 |
| Habitat:Maximum temperature | 0.707909 | 74 | 0.867 |
| Habitat:Minimum temperature | 0.023129 | 71 | 0.786 |
| Region:Altitude | 975.3777 | 66 | 0.001 |
| Region:Maximum temperature | 1.031212 | 61 | 0.157 |
| Region:Minimum temperature | 0.010138 | 56 | 0.696 |
| Sample site:Altitude | 1039.524 | 44 | 0.001 |
| Altitude:Maximum temperature | 6.41E-06 | 43 | 0.962 |
| Altitude:Minimum temperature | 6.33E-05 | 42 | 0.501 |
| Maximum temperature :Minimum temperature | 0.128122 | 41 | 0.876 |

### Influence of biological factors

Co-occurrence was common across the ash rot holes sampled, with 55.08 % of taxa combinations occurring only once or less (*n* = 18,403). Of the co-occurrences (*n* = 15,008), 10.9 % (*n* = 1,630) were considered non-random, with a large proportion occurring more often than expected (*n* = 1,429; 87.67 % of non-random co-occurrences). Uropodidae (Mesostigmata) had the most positive co-occurrences (taxa co-occurring more than expected; *n* = 46) followed by Acaridae (Oribatida) and Muscidae (Diptera; both *n* = 40). Arionidae (Gastropoda) had the most negative co-occurrences (*n* = 15) followed by Canthocamptidae (Copepoda; *n* = 11) and Thripidae (Thysanoptera; *n* = 10).

## Discussion

Ash rot holes are inhabited by a diverse entomofauna (almost 260 families), the community structure of which is driven by both biogeographical and tree-specific characteristics. Increasing ash veteran tree vulnerability makes sustaining these vital habitats and their associated invertebrate communities a conservation imperative. This study highlights the urgent need for integrated, cross-regional coordination of habitat management to maintain landscape-scale heterogeneity and rotting wood habitats to retain invertebrate diversity.

### Rot hole invertebrate community structure in veteran ash

The large diversity of families identified aligns with previously recorded rot hole inhabitants in other tree species (Irmler *et al*., 1996; Schauer *et al*., 2018; Cuff *et al*., 2021b; Schauer *et al*., 2025). Acari and Coleoptera were the most diverse groups, justifying them being the focus of most rot hole invertebrate research (Sverdrup-Thygeson *et al*., 2010; Müller *et al*., 2014; Taylor & Ranius, 2014; Pilskog *et al*., 2020; Cuff *et al*., 2021a; Lindman *et al*., 2023). Many of the other taxa identified have not been recorded in previous rot hole studies. Ants (Formicidae) were the sole Hymenoptera family in beech rot samples (Cuff *et al*., 2021b), yet the current study identified 22 Hymenoptera families, mostly comprising parasitoid wasps. Previous studies suggest that the presence of parasitoid wasps in rot hole communities is rare (Schauer *et al*., 2018, 2025; Cuff *et al*., 2021b), despite almost all insect orders and many non-insect arthropods being attacked by them (Fei *et al*., 2023). The high diversity of parasitoids within the present study may have been due to the larger sample size encompassing a greater spatial area across more habitat types than previous studies, rather the inherent biological differences between beech and ash rot holes.

Other groups that have only been recorded infrequently previously include mites, with only four mite species (all gall mites) previously listed as associated with ash (Mitchell *et al*. 2014; Littlewood *et al*. 2015); however, 69 families were recorded in the present study. The significant expansion of recorded ash-associated invertebrate families suggests an underestimation in past studies. Part of the challenge is that some taxa and some life stages (e.g., larvae) are difficult to identify morphologically. Many insects were represented by larval stages, perhaps due to the time of sampling, with many organisms beginning to overwinter. Molecular methods could be helpful (Sire *et al*., 2025), although incomplete barcode reference databases are a current limitation for many taxonomic groups (Cheng *et al*., 2023; Recuero et al. 2024; Sire *et al.,* 2025). Ecoacoustics, increasingly being applied to soil invertebrate monitoring (Metcalf *et al*., 2024; Robinson *et al*., 2024; Stroud *et al*., 2025), could also be used to determine arthropod diversity and activity (McAndrew *et al*., 2025) and in conjunction with other approaches, to detect the ecological interactions underway within these habitats (Dawson *et al*., 2026). Such approaches may, however, be less appropriate for monitoring smaller invertebrates such as mites (Schauer *et al*. 2025).

### The influence of tree-scale factors on rot hole invertebrate community structure

Several of the drivers of rot hole community structure identified in the present study align with the findings of previous studies focused on different tree species (Quinto *et al*., 2014; Schauer *et al*., 2018; Cuff *et al*., 2021b; Della Rocca *et al*., 2023). Wood rot water content and density were dominant tree-scale drivers of community structure in the present study and in beech (Cuff *et al*., 2021b). In most studies, however, there was a negative correlation between rot water content and saproxylic invertebrate diversity (Henneberg *et al*., 2021; Hernández-Corral *et al*., 2021; but see Ranius, 2002) although some taxa favour higher rot water content (Della Rocca *et al*., 2023). In the current study, only one insect family (Apidae) was associated with high water content, despite the association of many Diptera larvae with aquatic habitats (e.g., Ceratopogonidae, Chironomidae, Syrphidae). The positive correlation between the northness direction of the rot hole and the water content indicates a potential impact of solar radiation on water content; investigating both temperature and sun exposure of rot hole may identify whether water content was reliant upon the direction or driven by other factors.

As for other tree species (Ranius, 2002; Sverdrup-Thygeson *et al*., 2010; Quinto *et al*. 2014; Buse *et al*. 2016; Cuff *et al*. 2021b; Lindman *et al*. 2023), other ash tree characteristics (e.g., tree circumference and the height of the rot hole) also drove invertebrate community structure. Tree circumference is likely linked to microclimatic stability (Gouix *et al*., 2015; Lindman et al., 2023) and tree age, and these will, in turn, impact invertebrate communities (Stokland *et al*., 2012). Likewise rot hole opening size is likely to influence internal microclimate (Schauer *et al*., 2018). The height of the rot hole on the tree was also related to the abundance of many families, with lower openings likely accommodating more opportunistic soil dwelling taxa and therefore more closely resembling soil invertebrate communities than those higher up the trunk (Taylor & Ranius, 2014). Invertebrate communities in rot holes at different heights require further research given the vertical stratification of arthropod communities found in other parts of trees (Beaulieu *et al*., 2010; Madji *et al*., 2025).

### The influence of biogeography on rot hole invertebrate community structure

Biogeographical factors influence rot hole invertebrate communities, often interacting with tree-scale characteristics. Maximum temperature was correlated with all tree-scale characteristics other than rot hole height and its opening direction, which depend, at least partly, on external damage at some point in the past. Habitats type also affects tree development (Oktavia *et al*., 2022) and environmental conditions in rot holes (e.g., sunlight exposure and elevated temperatures; Sebek *et al*., 2016; Parmain & Bouget, 2018) will differ with tree density.

Most previous research has focused on non-urban woodland (e.g., Ranius, 2002; Schauer *et al*., 2018) despite the ecological importance of urban veteran trees (Gossmann *et al*., 2025; Parsons *et al*., 2025), their vulnerability to urban development (Parsons *et al*., 2023; Béland *et al*., 2025) and associated public safety concerns (Carpaneto *et al*., 2010). The current study revealed differences between habitat type, with a high diversity of invertebrates in urban woodland, and communities distinct from those in other woodland sites. These differences are possibly due to variation in conditions such as higher temperatures associated with urban heat islands (Deilami *et al*., 2018), and greater isolation due to habitat fragmentation (Nolan *et al*., 2022).

Parkland and farmland habitat also host many veteran ash (Perry, 2013; Mitchel *et al*., 2021; Ancient Tree Inventory, 2023) yet trees in these habitats tend to be overlooked in rot hole research. Parkland and farmland trees can provide refugia for many novel and rare species (Alexander, 1995; Cuff *et al*., 2021a; Wainhouse *et al*., 2024). The present study highlighted that their associated rot holes harbour unique and diverse communities. It also detected females of a potentially novel species belonging to the *Puliciphora* (Diptera: Phoridae) genus, males of which had been previously discovered from rot holes at one site (Dinefwr, Carmarthenshire, Wales; Welch *et al*., 2024). These females were required to confirm its identity and showed that it did not match any of the species in the Disney Collection of World Phoridae (Museum of Zoology, Cambridge University).

Many veteran trees within agricultural fields and margins are highly threatened (Hartel *et al*., 2017; Lindenmayer & Laurance, 2017; Prevedello *et al*., 2018) yet not only are they important ecologically but also from a farming perspective. Rot holes provide refugia for biocontrol agents of crop pests in farmland, as shown in the present study by the high frequency of natural enemies (e.g., parasitoid wasps), which were more abundant in rot holes in farmland than in woodland. Other crop pest natural enemies, such as the Trombidiformes mites (e.g., Trombidiidae; Knapp *et al*., 2018) were also more abundant in rot holes in farmland than in other habitat types. Rot holes may play an important role in maintaining overwintering invertebrate populations (Hassanpour *et al*., 2021). Whether these known natural enemies found in rot holes in the present study are attacking pests or saproxylic invertebrates, or simply overwintering, requires further empirical investigation as the presence of rot holes within agricultural systems may encourage the persistence of these and other biocontrol agents (Athey *et al*., 2016; Cuff *et al*., 2025).

### Future questions in rot hole invertebrate research

The present study has provided insight into community composition and the effects of some large scale biotic and abiotic factors on this. Investigation of the vertical stratification of invertebrates in rot holes, rot volume, wood rot fungal community and chemical composition, are needed to deepen understanding of these communities. Rot volume, for example, is a significant driver of community composition (Schauer *et al*., 2025) and could be measured alongside the overall volume and topography of rot holes using recent scanning approaches (Visick & Ratnieks, 2025).

Assessing ecological networks, including trophic and non-trophic interactions, can provide insight into the resilience of communities to perturbation (Pocock *et al*., 2012). The few examples where this approach has been applied to rot holes are, however, taxonomically restricted (Sánchez-Galván *et al*., 2018; Wetherbee *et al*., 2022), and the inability to observe the cryptic interactions in rot holes directly limits traditional approaches to their study. Molecular approaches (e.g., dietary metabarcoding; Pompanon *et al*., 2012), stable isotope analysis (Majdi *et al*., 2025) or the merging of multiple methodologies (Cuff *et al*., 2023; Dawson *et al*., 2026) could enable construction of trophic networks for rot hole communities (Windsor et al., 2023). Habitat connectivity analysis is also important (Pilshog et al. 2016; Sverdrup-Thygeson et al. 2017; Mestre et al. 2018). Further, to date, studies have been at single or just a few timepoints but to understand and model resilience of rot hole habitats and their whole communities under climate change scenarios requires long term monitoring.

## Conclusion

A vast diversity of invertebrates (259 families within the present study, and likely many more) inhabit ash rot holes, highlighting the need to conserve these communities across landscapes to sustain biodiversity. These communities are influenced by a plethora of biological and physical factors, including structural characteristics of both rot holes and their host trees, and by broader biogeographical factors, emphasising the importance of safeguarding the heterogeneity of these habitats. By bridging local- and landscape-scale drivers of community structure, this study provides a uniquely integrative understanding of the dynamics underpinning rot hole communities. Nevertheless, more research is needed into other tree species and biogeographical contexts and expanding our understanding through approaches like network ecology will further enhance our appreciation and understanding of these keystone woodland microhabitats.

## Acknowledgements

We thank Natural England for funding, Matt Wainhouse for his support, and Ed Woolley for help with visiting field sites. For permitting sampling to be conducted on sites, we would like to acknowledge: National Trust (Attingham Park, Dinefwr, Horner Woods, Sherborne Park Estate); Wildlife Trust of South & West Wales (Coed Dyrysiog, Coed Maidie B Goddard, Coed Penglanowen, Pwll y Wrach); Trustees of Dinder Estate (Dinder Woods); Friends of Forrest Farm, Cardiff Council (Forest Farm Country Park); Hayes Farm; Dr Rob Thomas, (Coed-ty-Llwyd Woods); Newcourt Farm; and Gloucester City Council (Robinswood Hill Country Park). For the purpose of open access, the author has applied a Creative Commons Attribution (CC BY) licence to any Author Accepted Manuscript version arising from this submission.

## Conflict of Interest

No conflicts of interest relevant to this manuscript.

## Author Contributions

WD: conceptualisation, sample collection, identification, data analysis and curation, writing – original draft; writing and editing. THJ: conceptualisation, supervision, editing. JPC: data analysis, editing. MS: identification, editing. LB: obtained funding, conceptualisation, supervision, editing.

## Data Availability

Data will be made available via Dryad at revision stage.

## Supplementary Materials

**Table S1.** Identification resources used throughout morphological identification to family level.

| Identification Resource | Taxon |
| --- | --- |
| Hopkin, S. 2014. *A key to the woodlice of Britain and Ireland*. 1st ed. Telford, UK: FSC Publications. | Woodlice |
| Unwin, D.M., 2001. *A key to the families of British Bugs (Insecta, Hemiptera)*. FSC Publications. | Hemiptera |
| Roberts, 1996. *Spiders: Britain and Northern Europe*. Harper Collins Publishers, Hammersmith, London.<br>Bee, L., Oxford, G. and Smith, H., 2017. Britain's spiders: a field guide. In *Britain's Spiders*. Princeton University Press. | Araneae |
| Unwin, D.M. 1988. *A key to the families of British coleoptera (and Strepsiptera)*. Williton, Somerset, FSC Publications.<br>Hammond, P.M., Marshall, J.E., Cox, M.L., Jessop, L., Garner, B.H. and Barclay, M.V.L. 2019. *British Coleoptera Larvae: A Guide to the Families and Major Subfamilies*. London, Royal Entomological Society. | Coleoptera |
| Evans, G.O. and Browning, E. 1954. *Pseudoscorpiones, with* | Pseudoscorpiones |
| keys to the species. London, Linnean Society of London.<br>Legg, G. and Farr-Cox, F. 2016. *Illustrated Key to the British False Scorpions:(pseudoscorpions)*. FSC Publications. |  |
| Gauld, I. and Bolton, B. 1988. *The Hymenoptera*. Oxford university press. | Hymenoptera |
| Barber, A.D., 2008. *Key to the identification of British centipedes*. FSC. | Centipedes |
| Smith, K.G.V. 1989. *An introduction to the immature stages of British flies: Diptera larvae, with notes on eggs, puparia and pupae*. 1st ed. London, UK: Royal Entomological Society of London.<br>Ball, S. 2008. *Introduction to the families of British Diptera*. Dipterists Forum. | Diptera |
| Shepherd, M. 2024. *A Key to the Soil Mites of Britain and Ireland*. [in prep] | Acari |
| Hopkin, S.P. 2007. *A key to the Collembola (springtails) of Britain and Ireland*. FSC Publications. | Collembola |
| Richards, P. 2010. *Guide to harvestmen of the British Isles*. FSC Publications. | Opiliones |
| Cameron, R. 2008. *Land Snails in the British Isles*. FSC Publications.<br>Rowson, B., Turner, J., Anderson, R., and Symondson, B. 2014. *Slugs of Britain and Ireland*. FSC Publications. | Gastropoda |
| Whitaker, A. 2007. *Fleas (Siphonaptera)*. *Handbooks for the Identification of British Insects Vol 1 (16)*. Royal Entomology Society. | Siphonaptera |
| New, T.R. 2005. *Psocids Psocoptera (Booklice and barklice)*. *Handbooks for the Identification of British Insects Vol 1 (7)*. Royal Entomology Society. | Psocoptera |
| Chinery, M. 2007. *Insects of Britain and Western Europe, domino field guide*. 2<sup>nd</sup> edn. A & C Black Publishers Ltd, London.<br>Tilling, S.M., 2014. *A key to the major groups of British terrestrial invertebrates*. Field Studies Council.<br>Mikes Insect Keys. 2023. *Mikes Insect Keys*. Basingstoke, North Hampshire [accessed at https://sites.google.com/view/mikes-insect-keys/mikes-insect-keys] | General Invertebrates |

**Table S2.**
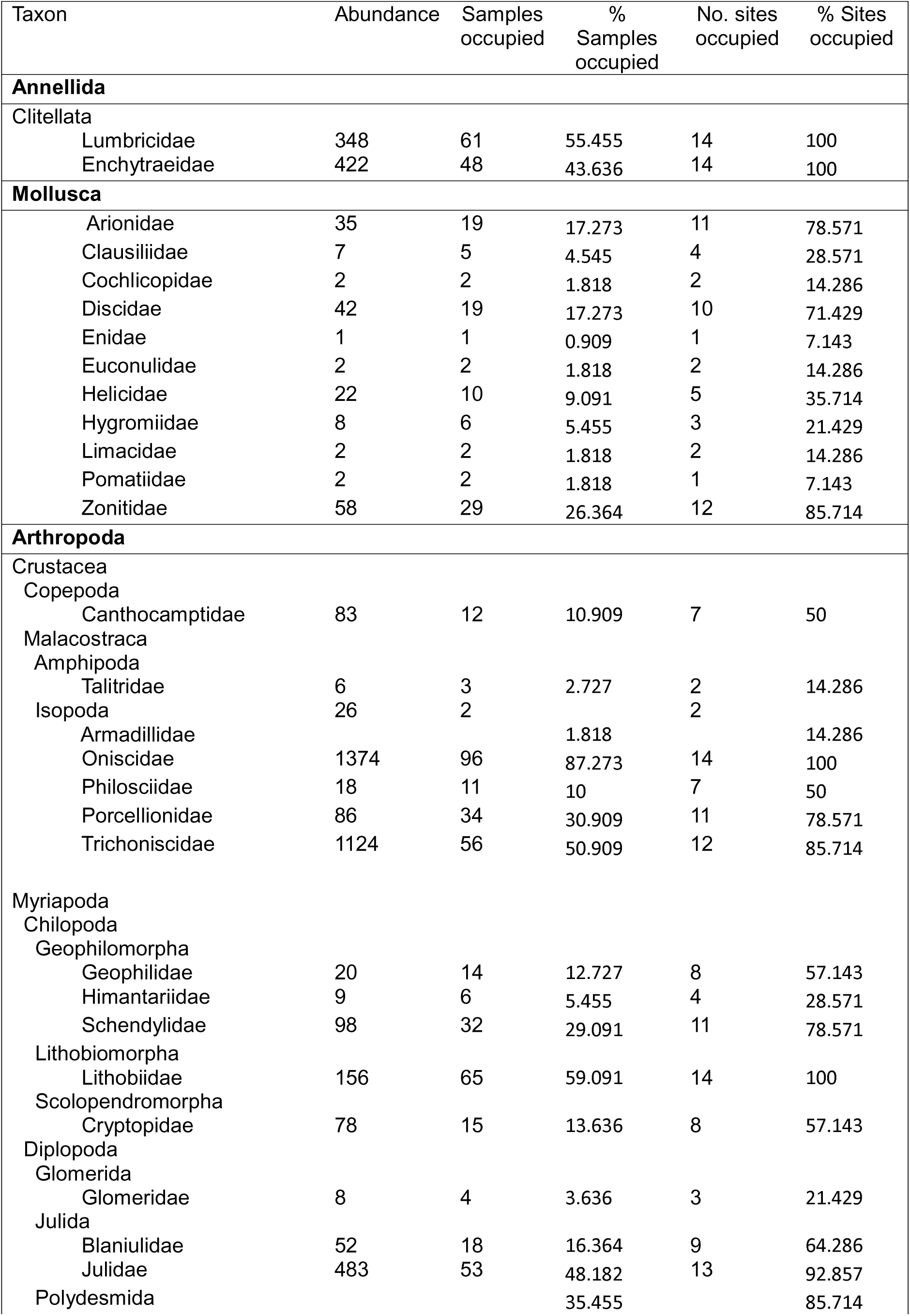
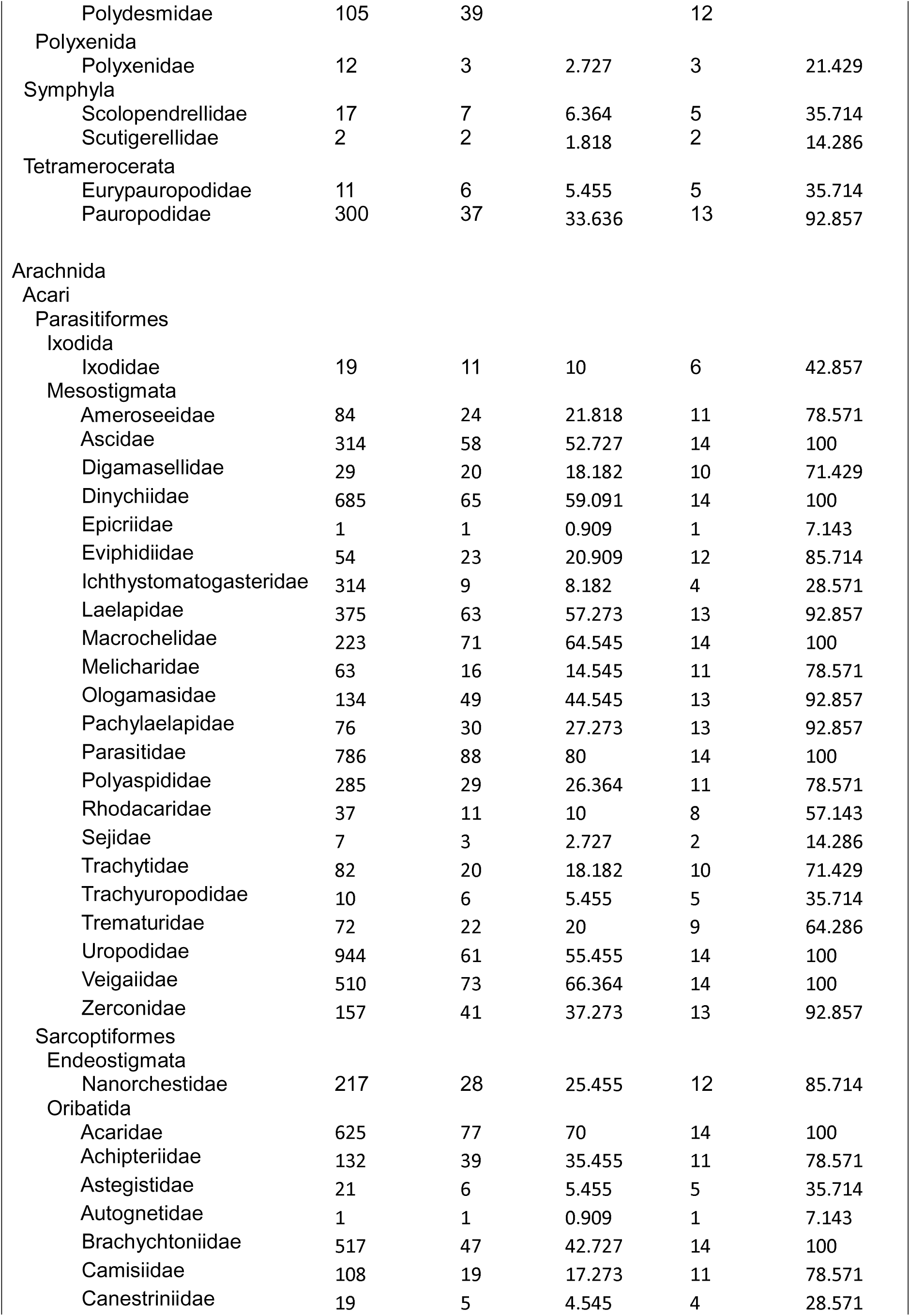
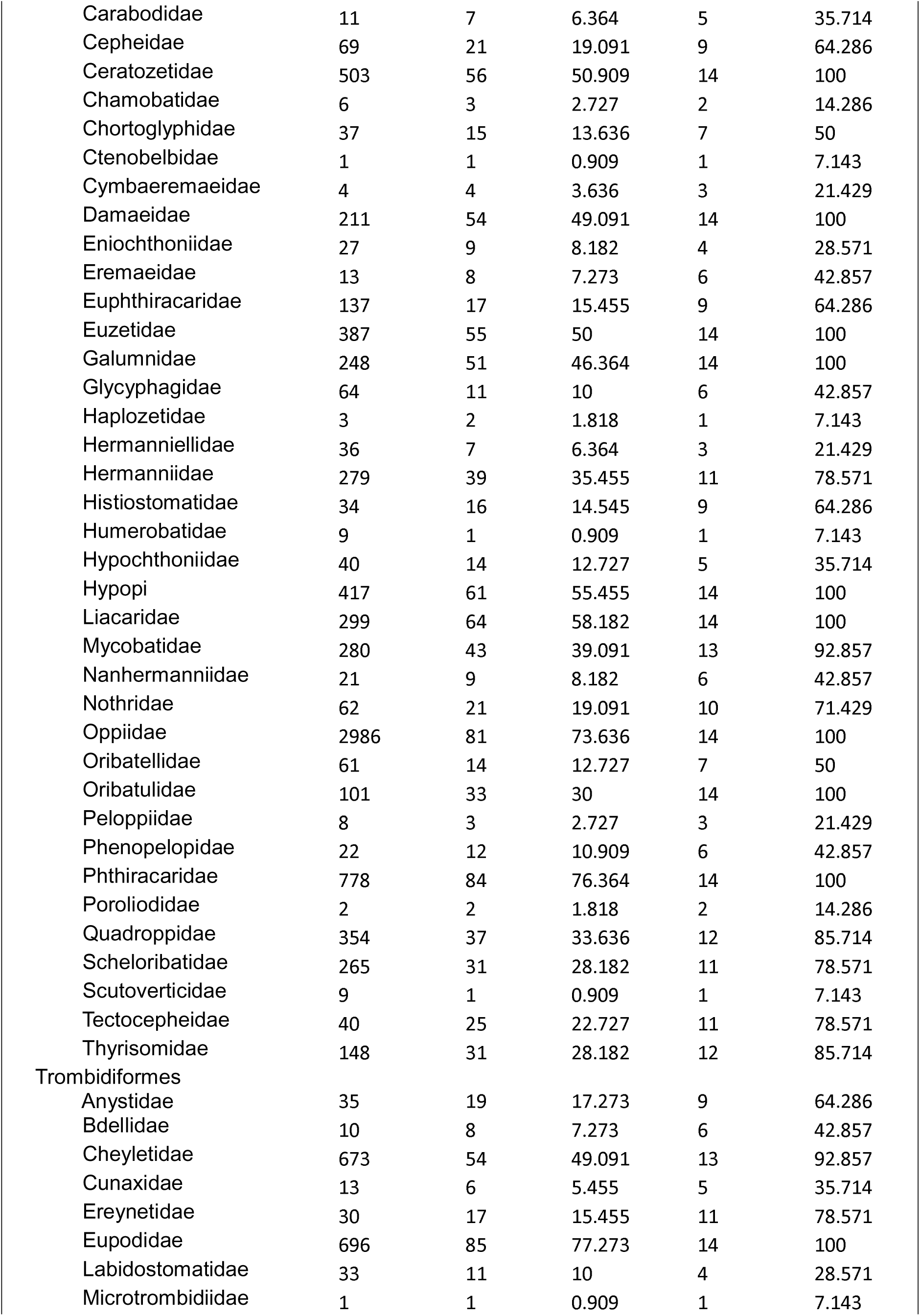
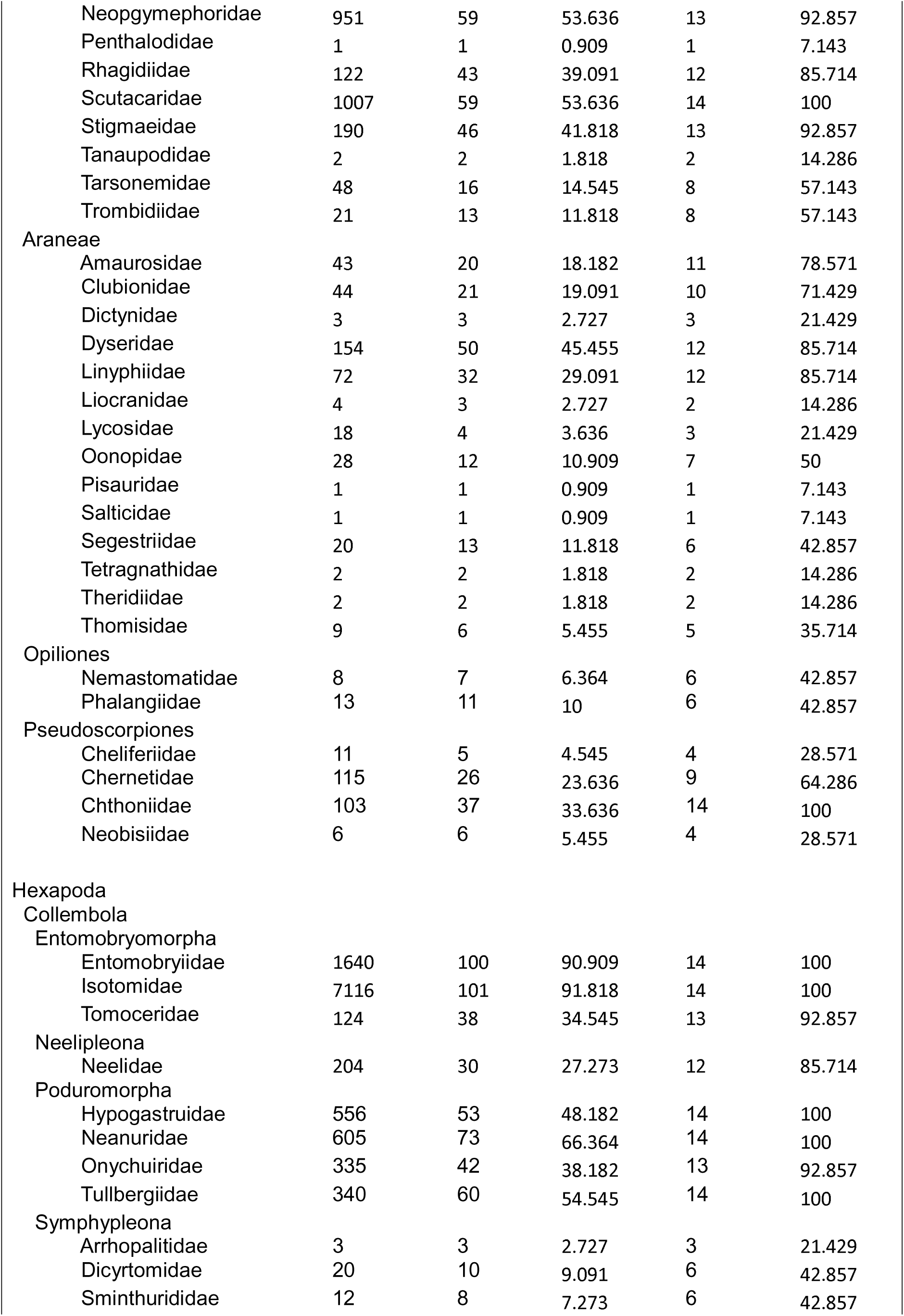
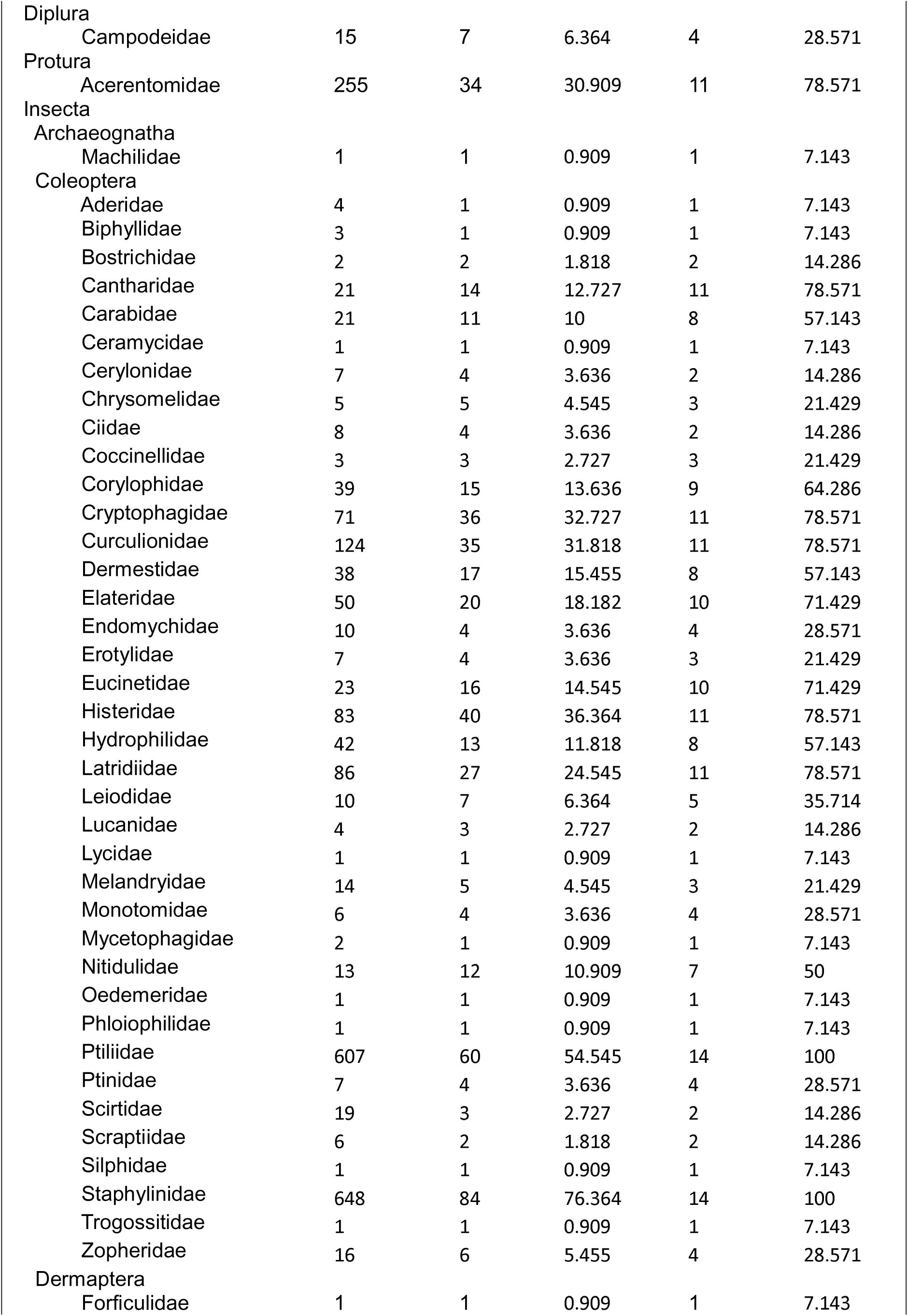
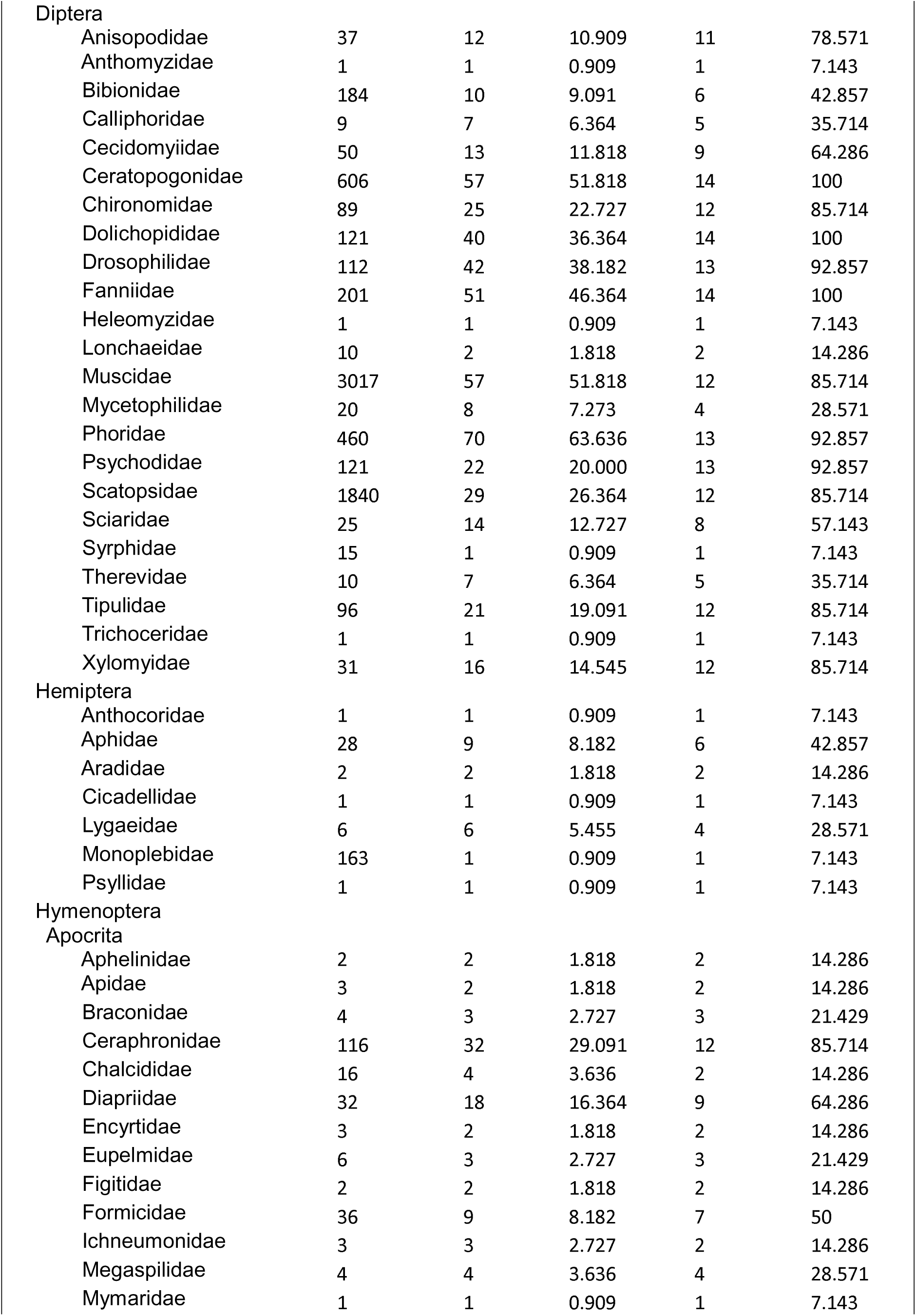
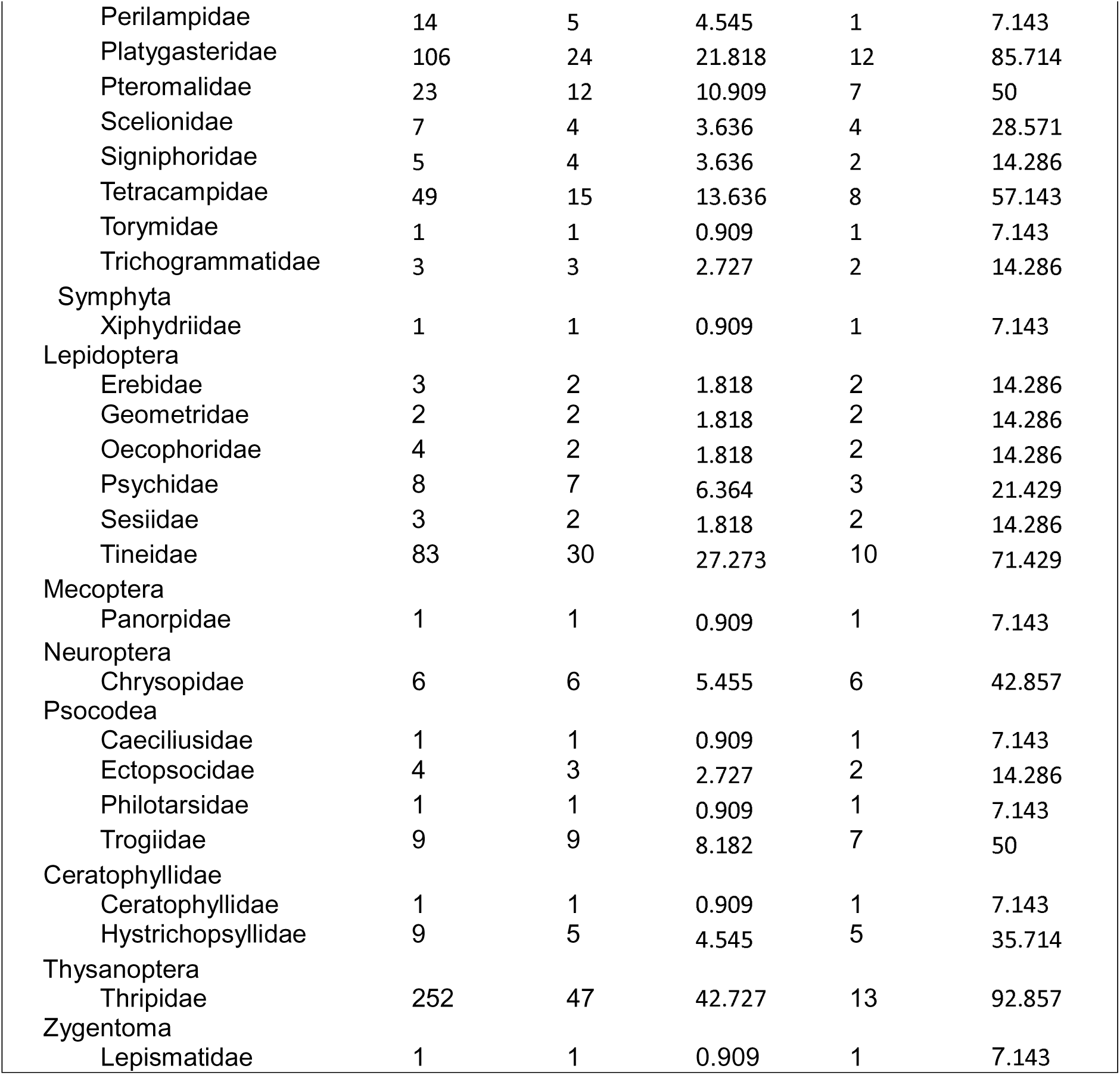
Summary of taxa identified. . All families that were identified are given in their rough taxonomic groups with their abundance across all samples, the number and percentage of samples the taxa occur within, and the number and percentage of regions they occur within.

| Taxon | Abundance | Samples occupied | % Samples occupied | No. sites occupied | % Sites occupied |
| --- | --- | --- | --- | --- | --- |
| **Annelida** |  |  |  |  |  |
| Clitellata |  |  |  |  |  |
| Lumbricidae | 348 | 61 | 55.455 | 14 | 100 |
| Enchytraeidae | 422 | 48 | 43.636 | 14 | 100 |
| **Mollusca** |  |  |  |  |  |
| Arionidae | 35 | 19 | 17.273 | 11 | 78.571 |
| Clausiliidae | 7 | 5 | 4.545 | 4 | 28.571 |
| Cochlicopidae | 2 | 2 | 1.818 | 2 | 14.286 |
| Discidae | 42 | 19 | 17.273 | 10 | 71.429 |
| Enidae | 1 | 1 | 0.909 | 1 | 7.143 |
| Euconulidae | 2 | 2 | 1.818 | 2 | 14.286 |
| Helicidae | 22 | 10 | 9.091 | 5 | 35.714 |
| Hygromiidae | 8 | 6 | 5.455 | 3 | 21.429 |
| Limacidae | 2 | 2 | 1.818 | 2 | 14.286 |
| Pomatiidae | 2 | 2 | 1.818 | 1 | 7.143 |
| Zonitidae | 58 | 29 | 26.364 | 12 | 85.714 |
| **Arthropoda** |  |  |  |  |  |
| Crustacea |  |  |  |  |  |
| Copepoda |  |  |  |  |  |
| Canthocamptidae | 83 | 12 | 10.909 | 7 | 50 |
| Malacostraca |  |  |  |  |  |
| Amphipoda |  |  |  |  |  |
| Talitridae | 6 | 3 | 2.727 | 2 | 14.286 |
| Isopoda | 26 | 2 |  | 2 |  |
| Armadillidae |  |  | 1.818 |  | 14.286 |
| Oniscidae | 1374 | 96 | 87.273 | 14 | 100 |
| Philosciidae | 18 | 11 | 10 | 7 | 50 |
| Porcellionidae | 86 | 34 | 30.909 | 11 | 78.571 |
| Trichoniscidae | 1124 | 56 | 50.909 | 12 | 85.714 |
| Myriapoda |  |  |  |  |  |
| Chilopoda |  |  |  |  |  |
| Geophilomorpha |  |  |  |  |  |
| Geophilidae | 20 | 14 | 12.727 | 8 | 57.143 |
| Himantariidae | 9 | 6 | 5.455 | 4 | 28.571 |
| Schendylidae | 98 | 32 | 29.091 | 11 | 78.571 |
| Lithobiomorpha |  |  |  |  |  |
| Lithobiidae | 156 | 65 | 59.091 | 14 | 100 |
| Scolopendromorpha |  |  |  |  |  |
| Cryptopidae | 78 | 15 | 13.636 | 8 | 57.143 |
| Diplopoda |  |  |  |  |  |
| Glomerida |  |  |  |  |  |
| Glomeridae | 8 | 4 | 3.636 | 3 | 21.429 |
| Julida |  |  |  |  |  |
| Blaniulidae | 52 | 18 | 16.364 | 9 | 64.286 |
| Julidae | 483 | 53 | 48.182 | 13 | 92.857 |
| Polydesmida |  |  | 35.455 |  | 85.714 |
| Polydesmidae | 105 | 39 |  | 12 |  |
| Polyxenida |  |  |  |  |  |
| Polyxenidae | 12 | 3 | 2.727 | 3 | 21.429 |
| Symphyla |  |  |  |  |  |
| Scolopendrellidae | 17 | 7 | 6.364 | 5 | 35.714 |
| Scutigerellidae | 2 | 2 | 1.818 | 2 | 14.286 |
| Tetramerocerata |  |  |  |  |  |
| Eurypauropodidae | 11 | 6 | 5.455 | 5 | 35.714 |
| Pauropodidae | 300 | 37 | 33.636 | 13 | 92.857 |
| Arachnida |  |  |  |  |  |
| Acari |  |  |  |  |  |
| Parasitiformes |  |  |  |  |  |
| Ixodida |  |  |  |  |  |
| Ixodidae | 19 | 11 | 10 | 6 | 42.857 |
| Mesostigmata |  |  |  |  |  |
| Ameroseeidae | 84 | 24 | 21.818 | 11 | 78.571 |
| Ascidae | 314 | 58 | 52.727 | 14 | 100 |
| Digamasellidae | 29 | 20 | 18.182 | 10 | 71.429 |
| Dinychiidae | 685 | 65 | 59.091 | 14 | 100 |
| Epicriidae | 1 | 1 | 0.909 | 1 | 7.143 |
| Eviphidiidae | 54 | 23 | 20.909 | 12 | 85.714 |
| Ichthystomatogasteridae | 314 | 9 | 8.182 | 4 | 28.571 |
| Laelapidae | 375 | 63 | 57.273 | 13 | 92.857 |
| Macrochelidae | 223 | 71 | 64.545 | 14 | 100 |
| Melicharidae | 63 | 16 | 14.545 | 11 | 78.571 |
| Ologamasidae | 134 | 49 | 44.545 | 13 | 92.857 |
| Pachylaelapidae | 76 | 30 | 27.273 | 13 | 92.857 |
| Parasitidae | 786 | 88 | 80 | 14 | 100 |
| Polyaspididae | 285 | 29 | 26.364 | 11 | 78.571 |
| Rhodacaridae | 37 | 11 | 10 | 8 | 57.143 |
| Sejidae | 7 | 3 | 2.727 | 2 | 14.286 |
| Trachytidae | 82 | 20 | 18.182 | 10 | 71.429 |
| Trachyuropodidae | 10 | 6 | 5.455 | 5 | 35.714 |
| Trematuridae | 72 | 22 | 20 | 9 | 64.286 |
| Uropodidae | 944 | 61 | 55.455 | 14 | 100 |
| Veigaiidae | 510 | 73 | 66.364 | 14 | 100 |
| Zerconidae | 157 | 41 | 37.273 | 13 | 92.857 |
| Sarcoptiformes |  |  |  |  |  |
| Endeostigmata |  |  |  |  |  |
| Nanorchestidae | 217 | 28 | 25.455 | 12 | 85.714 |
| Oribatida |  |  |  |  |  |
| Acaridae | 625 | 77 | 70 | 14 | 100 |
| Achipteriidae | 132 | 39 | 35.455 | 11 | 78.571 |
| Astegistidae | 21 | 6 | 5.455 | 5 | 35.714 |
| Autognetidae | 1 | 1 | 0.909 | 1 | 7.143 |
| Brachychtoniidae | 517 | 47 | 42.727 | 14 | 100 |
| Camisiidae | 108 | 19 | 17.273 | 11 | 78.571 |
| Canestriniidae | 19 | 5 | 4.545 | 4 | 28.571 |
| Carabodidae | 11 | 7 | 6.364 | 5 | 35.714 |
| Cepheidae | 69 | 21 | 19.091 | 9 | 64.286 |
| Ceratozetidae | 503 | 56 | 50.909 | 14 | 100 |
| Chamobatidae | 6 | 3 | 2.727 | 2 | 14.286 |
| Chortoglyphidae | 37 | 15 | 13.636 | 7 | 50 |
| Ctenobelbidae | 1 | 1 | 0.909 | 1 | 7.143 |
| Cymbaeremaeidae | 4 | 4 | 3.636 | 3 | 21.429 |
| Damaeidae | 211 | 54 | 49.091 | 14 | 100 |
| Eniochthoniidae | 27 | 9 | 8.182 | 4 | 28.571 |
| Eremaeidae | 13 | 8 | 7.273 | 6 | 42.857 |
| Euphthiracaridae | 137 | 17 | 15.455 | 9 | 64.286 |
| Euzetidae | 387 | 55 | 50 | 14 | 100 |
| Galumnidae | 248 | 51 | 46.364 | 14 | 100 |
| Glycyphagidae | 64 | 11 | 10 | 6 | 42.857 |
| Haplozetidae | 3 | 2 | 1.818 | 1 | 7.143 |
| Hermanniellidae | 36 | 7 | 6.364 | 3 | 21.429 |
| Hermanniidae | 279 | 39 | 35.455 | 11 | 78.571 |
| Histiostomatidae | 34 | 16 | 14.545 | 9 | 64.286 |
| Humerobatidae | 9 | 1 | 0.909 | 1 | 7.143 |
| Hypochthoniidae | 40 | 14 | 12.727 | 5 | 35.714 |
| Hypopi | 417 | 61 | 55.455 | 14 | 100 |
| Liacaridae | 299 | 64 | 58.182 | 14 | 100 |
| Mycobatidae | 280 | 43 | 39.091 | 13 | 92.857 |
| Nanhermanniidae | 21 | 9 | 8.182 | 6 | 42.857 |
| Nothridae | 62 | 21 | 19.091 | 10 | 71.429 |
| Oppiidae | 2986 | 81 | 73.636 | 14 | 100 |
| Oribatellidae | 61 | 14 | 12.727 | 7 | 50 |
| Oribatulidae | 101 | 33 | 30 | 14 | 100 |
| Peloppiidae | 8 | 3 | 2.727 | 3 | 21.429 |
| Phenopelopidae | 22 | 12 | 10.909 | 6 | 42.857 |
| Phthiracaridae | 778 | 84 | 76.364 | 14 | 100 |
| Poroliodidae | 2 | 2 | 1.818 | 2 | 14.286 |
| Quadroppidae | 354 | 37 | 33.636 | 12 | 85.714 |
| Scheloribatidae | 265 | 31 | 28.182 | 11 | 78.571 |
| Scutoverticidae | 9 | 1 | 0.909 | 1 | 7.143 |
| Tectocepheidae | 40 | 25 | 22.727 | 11 | 78.571 |
| Thyrisomidae | 148 | 31 | 28.182 | 12 | 85.714 |
| Trombidiformes |  |  |  |  |  |
| Anystidae | 35 | 19 | 17.273 | 9 | 64.286 |
| Bdellidae | 10 | 8 | 7.273 | 6 | 42.857 |
| Cheyletidae | 673 | 54 | 49.091 | 13 | 92.857 |
| Cunaxidae | 13 | 6 | 5.455 | 5 | 35.714 |
| Ereynetidae | 30 | 17 | 15.455 | 11 | 78.571 |
| Eupodidae | 696 | 85 | 77.273 | 14 | 100 |
| Labidostomatidae | 33 | 11 | 10 | 4 | 28.571 |
| Microtrombidiidae | 1 | 1 | 0.909 | 1 | 7.143 |
| Neopgymephoridae | 951 | 59 | 53.636 | 13 | 92.857 |
| Penthalodidae | 1 | 1 | 0.909 | 1 | 7.143 |
| Rhagidiidae | 122 | 43 | 39.091 | 12 | 85.714 |
| Scutacaridae | 1007 | 59 | 53.636 | 14 | 100 |
| Stigmaeidae | 190 | 46 | 41.818 | 13 | 92.857 |
| Tanaupodidae | 2 | 2 | 1.818 | 2 | 14.286 |
| Tarsonemidae | 48 | 16 | 14.545 | 8 | 57.143 |
| Trombidiidae | 21 | 13 | 11.818 | 8 | 57.143 |
| Araneae |  |  |  |  |  |
| Amaurosidae | 43 | 20 | 18.182 | 11 | 78.571 |
| Clubionidae | 44 | 21 | 19.091 | 10 | 71.429 |
| Dictynidae | 3 | 3 | 2.727 | 3 | 21.429 |
| Dyseridae | 154 | 50 | 45.455 | 12 | 85.714 |
| Linyphiidae | 72 | 32 | 29.091 | 12 | 85.714 |
| Liocranidae | 4 | 3 | 2.727 | 2 | 14.286 |
| Lycosidae | 18 | 4 | 3.636 | 3 | 21.429 |
| Oonopidae | 28 | 12 | 10.909 | 7 | 50 |
| Pisauridae | 1 | 1 | 0.909 | 1 | 7.143 |
| Salticidae | 1 | 1 | 0.909 | 1 | 7.143 |
| Segestriidae | 20 | 13 | 11.818 | 6 | 42.857 |
| Tetragnathidae | 2 | 2 | 1.818 | 2 | 14.286 |
| Theridiidae | 2 | 2 | 1.818 | 2 | 14.286 |
| Thomisidae | 9 | 6 | 5.455 | 5 | 35.714 |
| Opiliones |  |  |  |  |  |
| Nemastomatidae | 8 | 7 | 6.364 | 6 | 42.857 |
| Phalangiidae | 13 | 11 | 10 | 6 | 42.857 |
| Pseudoscorpiones |  |  |  |  |  |
| Cheliferiidae | 11 | 5 | 4.545 | 4 | 28.571 |
| Chernetidae | 115 | 26 | 23.636 | 9 | 64.286 |
| Chthoniidae | 103 | 37 | 33.636 | 14 | 100 |
| Neobisiidae | 6 | 6 | 5.455 | 4 | 28.571 |
| Hexapoda |  |  |  |  |  |
| Collembola |  |  |  |  |  |
| Entomobryomorpha |  |  |  |  |  |
| Entomobryiidae | 1640 | 100 | 90.909 | 14 | 100 |
| Isotomidae | 7116 | 101 | 91.818 | 14 | 100 |
| Tomoceridae | 124 | 38 | 34.545 | 13 | 92.857 |
| Neelipleona |  |  |  |  |  |
| Neelidae | 204 | 30 | 27.273 | 12 | 85.714 |
| Poduromorpha |  |  |  |  |  |
| Hypogastruidae | 556 | 53 | 48.182 | 14 | 100 |
| Neanuridae | 605 | 73 | 66.364 | 14 | 100 |
| Onychiuridae | 335 | 42 | 38.182 | 13 | 92.857 |
| Tullbergiidae | 340 | 60 | 54.545 | 14 | 100 |
| Symphypleona |  |  |  |  |  |
| Arrhopalitidae | 3 | 3 | 2.727 | 3 | 21.429 |
| Dicyrtomidae | 20 | 10 | 9.091 | 6 | 42.857 |
| Sminthurididae | 12 | 8 | 7.273 | 6 | 42.857 |
| Diplura |  |  |  |  |  |
| Campodeidae | 15 | 7 | 6.364 | 4 | 28.571 |
| Protura |  |  |  |  |  |
| Acerentomidae | 255 | 34 | 30.909 | 11 | 78.571 |
| Insecta |  |  |  |  |  |
| Archaeognatha |  |  |  |  |  |
| Machilidae | 1 | 1 | 0.909 | 1 | 7.143 |
| Coleoptera |  |  |  |  |  |
| Aderidae | 4 | 1 | 0.909 | 1 | 7.143 |
| Biphyllidae | 3 | 1 | 0.909 | 1 | 7.143 |
| Bostrichidae | 2 | 2 | 1.818 | 2 | 14.286 |
| Cantharidae | 21 | 14 | 12.727 | 11 | 78.571 |
| Carabidae | 21 | 11 | 10 | 8 | 57.143 |
| Ceramycidae | 1 | 1 | 0.909 | 1 | 7.143 |
| Cerylonidae | 7 | 4 | 3.636 | 2 | 14.286 |
| Chrysomelidae | 5 | 5 | 4.545 | 3 | 21.429 |
| Ciidae | 8 | 4 | 3.636 | 2 | 14.286 |
| Coccinellidae | 3 | 3 | 2.727 | 3 | 21.429 |
| Corylophidae | 39 | 15 | 13.636 | 9 | 64.286 |
| Cryptophagidae | 71 | 36 | 32.727 | 11 | 78.571 |
| Curculionidae | 124 | 35 | 31.818 | 11 | 78.571 |
| Dermestidae | 38 | 17 | 15.455 | 8 | 57.143 |
| Elateridae | 50 | 20 | 18.182 | 10 | 71.429 |
| Endomychidae | 10 | 4 | 3.636 | 4 | 28.571 |
| Erotylidae | 7 | 4 | 3.636 | 3 | 21.429 |
| Eucinetidae | 23 | 16 | 14.545 | 10 | 71.429 |
| Histeridae | 83 | 40 | 36.364 | 11 | 78.571 |
| Hydrophilidae | 42 | 13 | 11.818 | 8 | 57.143 |
| Latridiidae | 86 | 27 | 24.545 | 11 | 78.571 |
| Leiodidae | 10 | 7 | 6.364 | 5 | 35.714 |
| Lucanidae | 4 | 3 | 2.727 | 2 | 14.286 |
| Lycidae | 1 | 1 | 0.909 | 1 | 7.143 |
| Melandryidae | 14 | 5 | 4.545 | 3 | 21.429 |
| Monotomidae | 6 | 4 | 3.636 | 4 | 28.571 |
| Mycetophagidae | 2 | 1 | 0.909 | 1 | 7.143 |
| Nitidulidae | 13 | 12 | 10.909 | 7 | 50 |
| Oedemeridae | 1 | 1 | 0.909 | 1 | 7.143 |
| Phloiophilidae | 1 | 1 | 0.909 | 1 | 7.143 |
| Ptiliidae | 607 | 60 | 54.545 | 14 | 100 |
| Ptinidae | 7 | 4 | 3.636 | 4 | 28.571 |
| Scirtidae | 19 | 3 | 2.727 | 2 | 14.286 |
| Scraptiidae | 6 | 2 | 1.818 | 2 | 14.286 |
| Silphidae | 1 | 1 | 0.909 | 1 | 7.143 |
| Staphylinidae | 648 | 84 | 76.364 | 14 | 100 |
| Trogossitidae | 1 | 1 | 0.909 | 1 | 7.143 |
| Zopheridae | 16 | 6 | 5.455 | 4 | 28.571 |
| Dermaptera |  |  |  |  |  |
| Forficulidae | 1 | 1 | 0.909 | 1 | 7.143 |
| Diptera |  |  |  |  |  |
| Anisopodidae | 37 | 12 | 10.909 | 11 | 78.571 |
| Anthomyzidae | 1 | 1 | 0.909 | 1 | 7.143 |
| Bibionidae | 184 | 10 | 9.091 | 6 | 42.857 |
| Calliphoridae | 9 | 7 | 6.364 | 5 | 35.714 |
| Cecidomyiidae | 50 | 13 | 11.818 | 9 | 64.286 |
| Ceratopogonidae | 606 | 57 | 51.818 | 14 | 100 |
| Chironomidae | 89 | 25 | 22.727 | 12 | 85.714 |
| Dolichopidae | 121 | 40 | 36.364 | 14 | 100 |
| Drosophilidae | 112 | 42 | 38.182 | 13 | 92.857 |
| Fanniidae | 201 | 51 | 46.364 | 14 | 100 |
| Heleomyzidae | 1 | 1 | 0.909 | 1 | 7.143 |
| Lonchaeidae | 10 | 2 | 1.818 | 2 | 14.286 |
| Muscidae | 3017 | 57 | 51.818 | 12 | 85.714 |
| Mycetophilidae | 20 | 8 | 7.273 | 4 | 28.571 |
| Phoridae | 460 | 70 | 63.636 | 13 | 92.857 |
| Psychodidae | 121 | 22 | 20.000 | 13 | 92.857 |
| Scatopsidae | 1840 | 29 | 26.364 | 12 | 85.714 |
| Sciaridae | 25 | 14 | 12.727 | 8 | 57.143 |
| Syrphidae | 15 | 1 | 0.909 | 1 | 7.143 |
| Therevidae | 10 | 7 | 6.364 | 5 | 35.714 |
| Tipulidae | 96 | 21 | 19.091 | 12 | 85.714 |
| Trichoceridae | 1 | 1 | 0.909 | 1 | 7.143 |
| Xylomyidae | 31 | 16 | 14.545 | 12 | 85.714 |
| Hemiptera |  |  |  |  |  |
| Anthocoridae | 1 | 1 | 0.909 | 1 | 7.143 |
| Aphidae | 28 | 9 | 8.182 | 6 | 42.857 |
| Aradidae | 2 | 2 | 1.818 | 2 | 14.286 |
| Cicadellidae | 1 | 1 | 0.909 | 1 | 7.143 |
| Lygaeidae | 6 | 6 | 5.455 | 4 | 28.571 |
| Monoplebidae | 163 | 1 | 0.909 | 1 | 7.143 |
| Psyllidae | 1 | 1 | 0.909 | 1 | 7.143 |
| Apocrita |  |  |  |  |  |
| Aphelinidae | 2 | 2 | 1.818 | 2 | 14.286 |
| Apidae | 3 | 2 | 1.818 | 2 | 14.286 |
| Braconidae | 4 | 3 | 2.727 | 3 | 21.429 |
| Ceraphronidae | 116 | 32 | 29.091 | 12 | 85.714 |
| Chalcididae | 16 | 4 | 3.636 | 2 | 14.286 |
| Diapriidae | 32 | 18 | 16.364 | 9 | 64.286 |
| Encyrtidae | 3 | 2 | 1.818 | 2 | 14.286 |
| Eupelmidae | 6 | 3 | 2.727 | 3 | 21.429 |
| Figitidae | 2 | 2 | 1.818 | 2 | 14.286 |
| Formicidae | 36 | 9 | 8.182 | 7 | 50 |
| Ichneumonidae | 3 | 3 | 2.727 | 2 | 14.286 |
| Megaspilidae | 4 | 4 | 3.636 | 4 | 28.571 |
| Mymaridae | 1 | 1 | 0.909 | 1 | 7.143 |
| Perilampidae | 14 | 5 | 4.545 | 1 | 7.143 |
| Platygasteridae | 106 | 24 | 21.818 | 12 | 85.714 |
| Pteromalidae | 23 | 12 | 10.909 | 7 | 50 |
| Scelionidae | 7 | 4 | 3.636 | 4 | 28.571 |
| Signiphoridae | 5 | 4 | 3.636 | 2 | 14.286 |
| Tetracampidae | 49 | 15 | 13.636 | 8 | 57.143 |
| Torymidae | 1 | 1 | 0.909 | 1 | 7.143 |
| Trichogrammatidae | 3 | 3 | 2.727 | 2 | 14.286 |
| Symphyta |  |  |  |  |  |
| Xiphydriidae | 1 | 1 | 0.909 | 1 | 7.143 |
| Lepidoptera |  |  |  |  |  |
| Erebidae | 3 | 2 | 1.818 | 2 | 14.286 |
| Geometridae | 2 | 2 | 1.818 | 2 | 14.286 |
| Oecophoridae | 4 | 2 | 1.818 | 2 | 14.286 |
| Psychidae | 8 | 7 | 6.364 | 3 | 21.429 |
| Sesiidae | 3 | 2 | 1.818 | 2 | 14.286 |
| Tineidae | 83 | 30 | 27.273 | 10 | 71.429 |
| Mecoptera |  |  |  |  |  |
| Panorpidae | 1 | 1 | 0.909 | 1 | 7.143 |
| Neuroptera |  |  |  |  |  |
| Chrysopidae | 6 | 6 | 5.455 | 6 | 42.857 |
| Psocodea |  |  |  |  |  |
| Caeciliusidae | 1 | 1 | 0.909 | 1 | 7.143 |
| Ectopsocidae | 4 | 3 | 2.727 | 2 | 14.286 |
| Philotarsidae | 1 | 1 | 0.909 | 1 | 7.143 |
| Trogiidae | 9 | 9 | 8.182 | 7 | 50 |
| Ceratophyllidae |  |  |  |  |  |
| Ceratophyllidae | 1 | 1 | 0.909 | 1 | 7.143 |
| Hystrihopsyllidae | 9 | 5 | 4.545 | 5 | 35.714 |
| Thysanoptera |  |  |  |  |  |
| Thripidae | 252 | 47 | 42.727 | 13 | 92.857 |
| Zygentoma |  |  |  |  |  |
| Lepismatidae | 1 | 1 | 0.909 | 1 | 7.143 |

**Table S3.** Summary of variation between tree-scale and biogeographical variables over site, region and habitat scales. The Kruskall-Wallis *X*^2^ values for investigating the variation in each variable across the different scales are provided.

| Variable | Sampling scale |  |  |
| --- | --- | --- | --- |
|  | Sample site (df = 13) | Region (df = 5) | Habitat (df = 3) |
| Circumference | 59.307 (p < 0.001) | 5.429 (p = 0.366) | 35.075 (p < 0.001) |
| Opening area | 35.458 (p < 0.001) | 9.321 (p = 0.097) | 10.695 (p = 0.014) |
| Height on tree | 21.724 (p = 0.060) | 8.412 (p = 0.135) | 4.727 (p = 0.193) |
| Northness of opening direction | 21.207 (p = 0.069) | 6.620 (p = 0.251) | 3.803 (p = 0.284) |
| Eastness of opening direction | 9.289 (p = 0.751) | 3.492 (p = 0.625) | 1.390 (p = 0.708) |
| Rot water content | 34.37 (p = 0.001) | 12.88 (p = 0.025) | 14.065 (p = 0.003) |
| Rot density | 32.051 (p = 0.002) | 18.422 (p = 0.002) | 13.147 (p = 0.004) |
| Altitude | 97.044 (p < 0.001) | 91.083 (p < 0.001) | 33.63 (p < 0.001) |
| Maximum temperature | 109 (p < 0.001) | 52.61 (p < 0.001) | 61.648 (p < 0.001) |
| Minimum temperature | 109 (p < 0.001) | 83.843 (p < 0.001) | 29.375 (p < 0.001) |
| Rainfall | 109 (p < 0.001) | 92.271 (p < 0.001) | 7.034 (p = 0.071) |

**Table S4.**
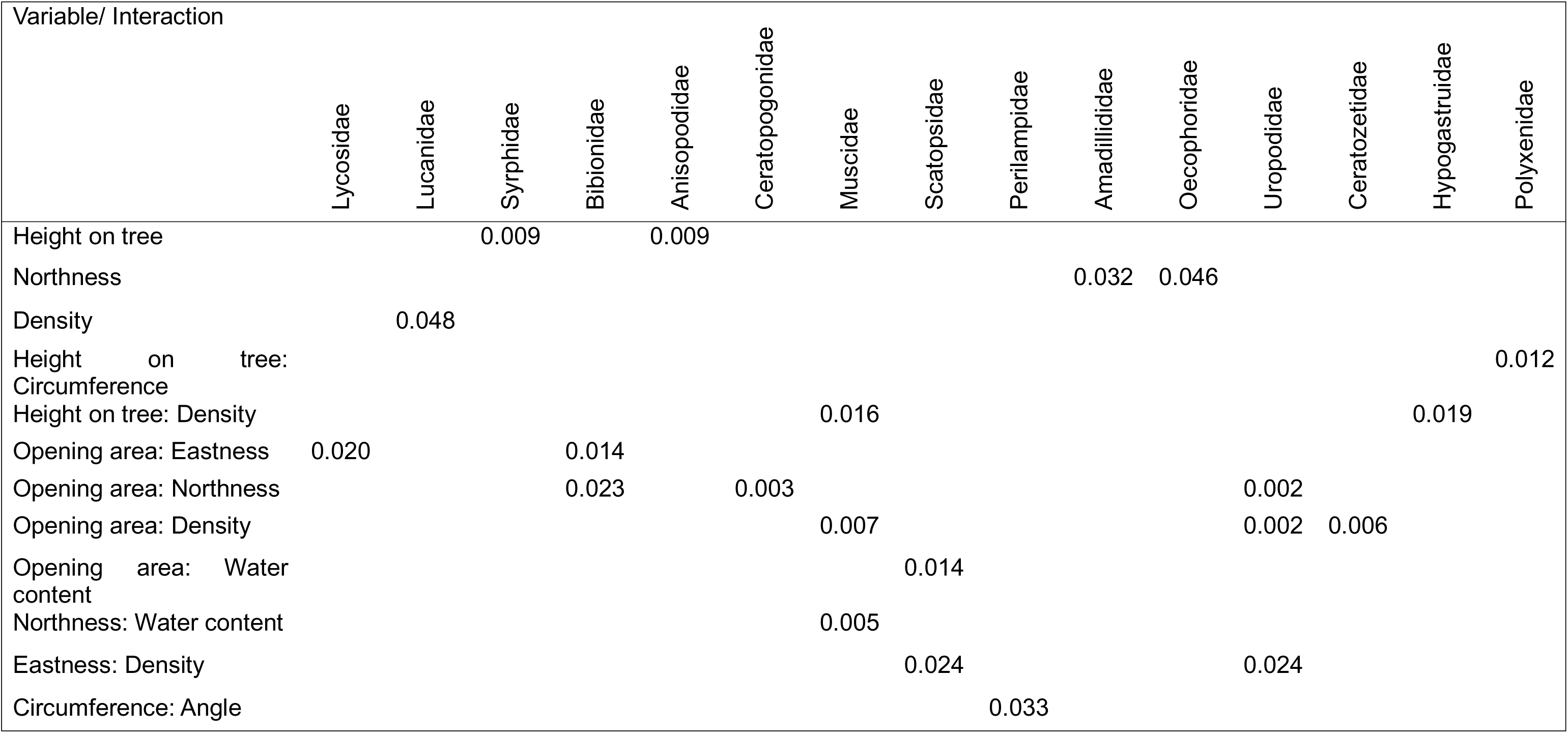
Significant influences of different tree-scale variables and their interactions upon individual rot hole taxa. The variables and interactions are given with the taxa they significantly influence (p < 0.05).

**Table S5.**
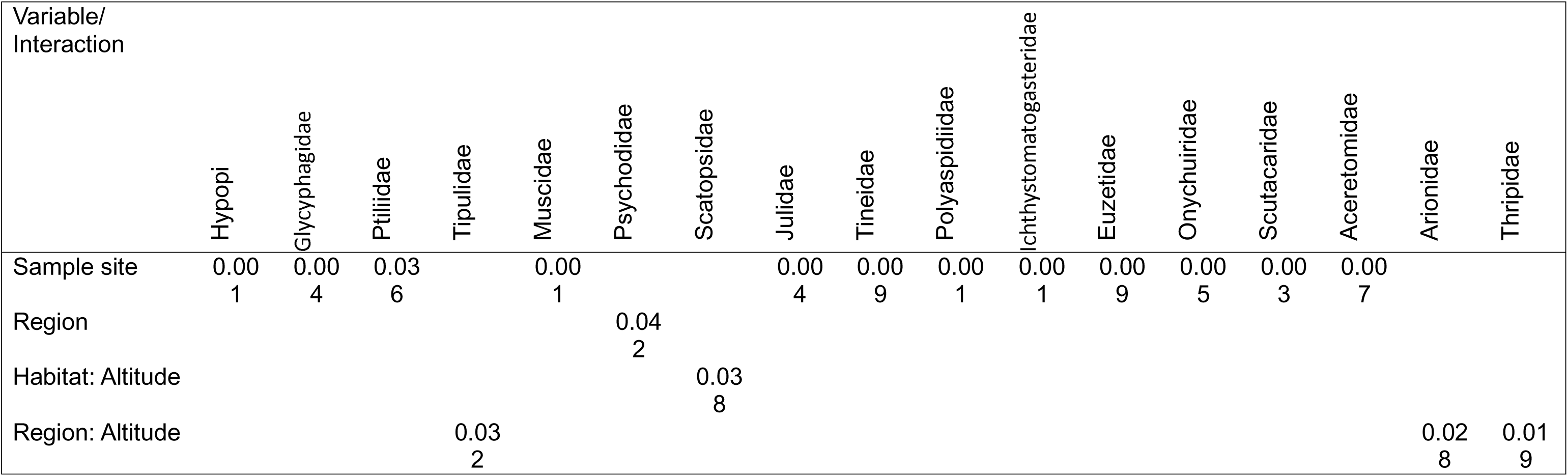
Significant influences of different biogeographic variables and their interactions upon individual rot hole taxa. The variables and interactions are given with the taxa they significantly influence (p < 0.05).

| Variable/<br>Interaction | Hypopi | Glycyphagidae | Ptiliidae | Tipulidae | Muscidae | Psychodidae | Scatopsidae | Julidae | Tineidae | Polyaspidiidae | Ichthyostomatogasteridae | Euzetidae | Onychuiridae | Scutacaridae | Aceretomidae | Arionidae | Thripidae |
| --- | --- | --- | --- | --- | --- | --- | --- | --- | --- | --- | --- | --- | --- | --- | --- | --- | --- |
| Sample site | 0.00<br>1 | 0.00<br>4 | 0.03<br>6 |  | 0.00<br>1 |  |  | 0.00<br>4 | 0.00<br>9 | 0.00<br>1 | 0.00<br>1 | 0.00<br>9 | 0.00<br>5 | 0.00<br>3 | 0.00<br>7 |  |  |
| Region |  |  |  |  |  | 0.04<br>2 |  |  |  |  |  |  |  |  |  |  |  |
| Habitat: Altitude |  |  |  |  |  |  | 0.03<br>8 |  |  |  |  |  |  |  |  |  |  |
| Region: Altitude |  |  |  | 0.03<br>2 |  |  |  |  |  |  |  |  |  |  |  | 0.02<br>8 | 0.01<br>9 |

**Figure S1.**
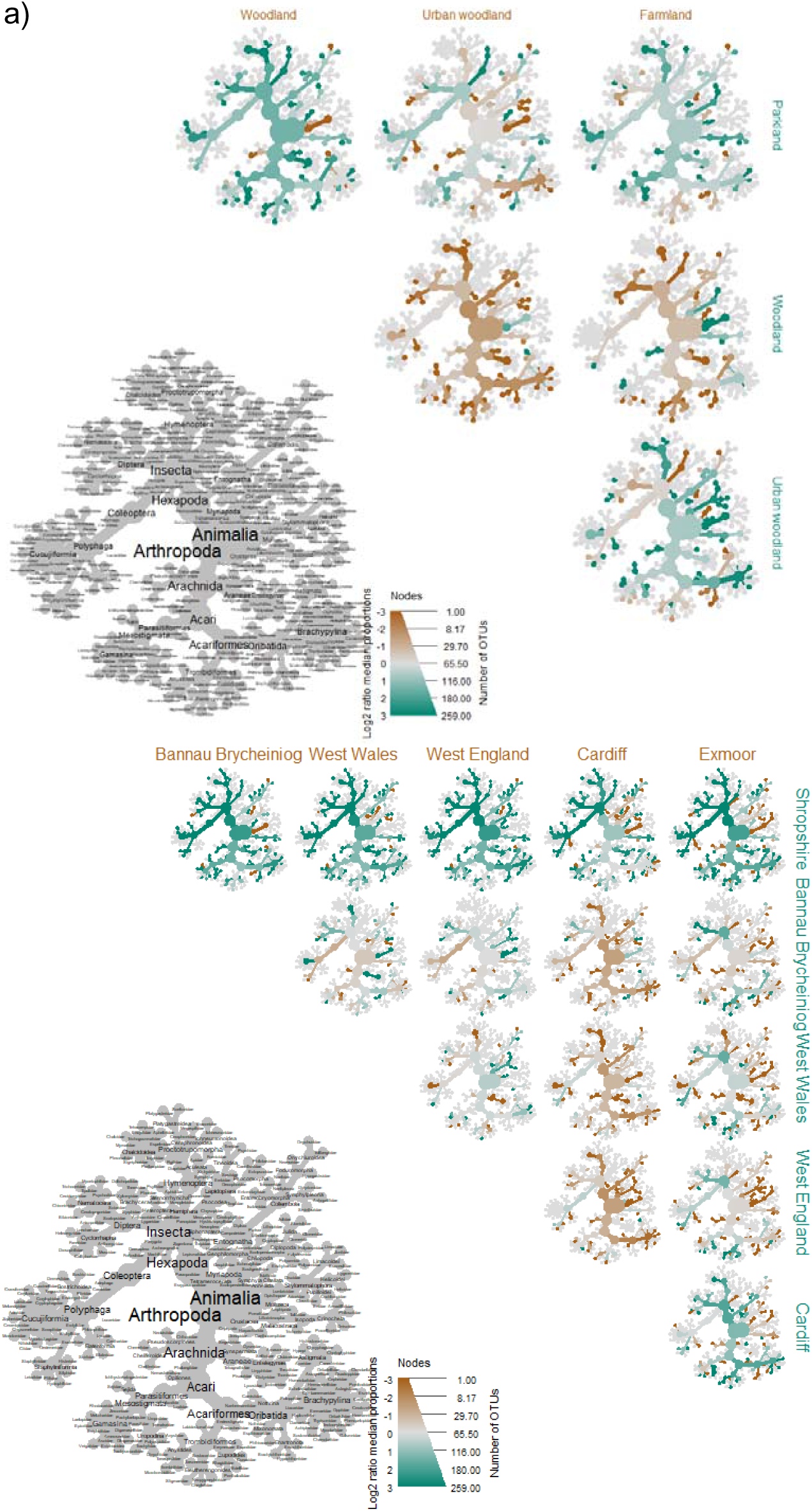
Community composition differences across different habitats and regions. The differences between the total community structures of each (a) habitat, and (b) region is plotted using ‘metacoder’. Taxa given in orange or green are more abundant in the corresponding geographical group. The total heat map plot corresponds to that in Figure.

## Notes

### Competing Interest Statement

The authors have declared no competing interest.

